# Structural basis of allosteric modulation of metabotropic glutamate receptor activation and desensitization

**DOI:** 10.1101/2023.08.13.552748

**Authors:** Alexa Strauss, Alberto J. Gonzalez-Hernandez, Joon Lee, Nohely Abreu, Purushotham Selvakumar, Leslie Salas-Estrada, Melanie Kristt, Dagan C. Marx, Kristen Gilliland, Bruce J. Melancon, Marta Filizola, Joel Meyerson, Joshua Levitz

**Affiliations:** Department of Biochemistry, Weill Cornell Medicine, New York, NY 10065, USA; Tri-Institutional Program in Chemical Biology, New York, NY 10065, USA; Department of Physiology and Biophysics, Weill Cornell Medicine, New York, NY 10065, USA; Department of Pharmacological Sciences, Icahn School of Medicine at Mount Sinai, New York, NY 10029, USA; Warren Center for Neuroscience Drug Discovery at Vanderbilt University, Vanderbilt University, Nashville, TN 37232, USA; Department of Pharmacology, Vanderbilt University School of Medicine, Nashville, TN 37232, USA; Department of Psychiatry, Weill Cornell Medicine, New York, NY 10065, USA

## Abstract

The metabotropic glutamate receptors (mGluRs) are neuromodulatory family C G protein coupled receptors which assemble as dimers and allosterically couple extracellular ligand binding domains (LBDs) to transmembrane domains (TMDs) to drive intracellular signaling. Pharmacologically, mGluRs can be targeted either at the LBDs by glutamate and synthetic orthosteric compounds or at the TMDs by allosteric modulators. Despite the potential of allosteric TMD-targeting compounds as therapeutics, an understanding of the functional and structural basis of their effects on mGluRs is limited. Here we use a battery of approaches to dissect the distinct functional and structural effects of orthosteric versus allosteric ligands. We find using electrophysiological and live cell imaging assays that both agonists and positive allosteric modulators (PAMs) can drive activation and desensitization of mGluRs. The effects of PAMs are pleiotropic, including both the ability to boost the maximal response to orthosteric agonists and to serve independently as desensitization-biased agonists across mGluR subtypes. Conformational sensors reveal PAM-driven inter-subunit re-arrangements at both the LBD and TMD. Motivated by this, we determine cryo-electron microscopy structures of mGluR3 in the presence of either an agonist or antagonist alone or in combination with a PAM. These structures reveal PAM-driven re-shaping of intra- and inter-subunit conformations and provide evidence for a rolling TMD dimer interface activation pathway that controls G protein and beta-arrestin coupling.

**Highlights:** -Agonists and PAMs drive mGluR activation, desensitization, and endocytosis

-PAMs are desensitization-biased and synergistic with agonists

-Four combinatorial ligand conditions reveal an ensemble of full-length mGluR structures with novel interfaces

-Activation and desensitization involve rolling TMD interfaces which are re-shaped by PAM

## Introduction

G protein coupled receptors (GPCRs) control a wide range of physiological and therapeutic processes by initiating intracellular signaling cascades via heterotrimeric G proteins^1,2^. Following activation, desensitization mechanisms tune GPCR signaling to maintain temporal precision and avoid excessive activation^3^. On the seconds to minutes time scale, acute desensitization can occur via G protein receptor kinases (GRKs), which bind to activated receptors to both sterically occlude G protein binding and sequester active G proteins^4–8^. GRKs also phosphorylate key intracellular residues of GPCRs to promote beta-arrestin (*β*-arr) coupling^9,10^ resulting in the recruitment of endocytic proteins to drive internalization and trafficking over the minutes to hour time scale^11^. A critical aspect of GPCR drug targeting is the ability of ligands to control both receptor activation and receptor desensitization. Different ligands for the same receptor are often able to stabilize distinct states within a receptor’s conformational landscape, which can lead to a relative bias towards modes of activation or desensitization^12–15^.

Determining the conformational basis of differential effects of ligands on GPCR activation and desensitization is a major challenge, particularly for the family C GPCRs. The metabotropic glutamate receptor (mGluRs) are prototypical family C GPCRs, which mediate synaptic neuromodulation and serve as prominent drug targets for a plethora of neurological and psychiatric disorders^16–18^. Structurally, mGluRs form constitutive dimers^19–22^ with each subunit containing a large extracellular domain consisting of a bi-lobed ligand binding domain (LBD) and a cysteine rich domain (CRD). The CRD connects to the canonical seven helix transmembrane domain (TMD) which is followed by a disordered C-terminal domain (CTD)^23^. The LBDs serve as the binding site for the native agonist glutamate as well as other synthetic orthosteric agonists and antagonists, while the TMDs both initiate transducer coupling and provide a binding site for synthetic positive (PAMs) and negative (NAMs) allosteric modulators^24,25^. PAMs are primarily defined by their ability to enhance the effect of agonists on G protein signaling^25–27^, but it has become increasingly clear that many mGluR PAMs can directly evoke activation in the full-length receptor in the absence of agonist^28–32^, thus serving as “ago-PAMs”. Understanding how PAMs control mGluR function and conformation is a critical challenge for both gaining a fundamental understanding of family C GPCRs and for harnessing the potential of PAMs as subtype-specific therapeutics^33,34^.

Over the last decade, structural and biophysical studies have shed light on the mechanisms of activation and allosteric modulation of mGluRs. At the dimeric LBDs, agonist-mediated activation is associated with substantial intrasubunit (i.e. clamshell closure) and intersubunit rearrangements that drive a common “closed-closed/active” state, as revealed by X-ray crystallography^36–38^ and Förster resonance energy transfer (FRET) studies^20,23,39–43^. Along with LBD rearrangements, full-length cryo-EM structures^30,38,44–48^ show an overall compaction of the CRD upon activation which is supported by recent single molecule FRET (smFRET) measurements^35,49^. At the TMD level, crystal structures of isolated TMDs^30,44,50,51^ as well as full length cryo-EM structures with different ligand conditions^30,38,44,46–48^ show variable inter-TMD interfaces, with strong evidence for agonist-driven stabilization of a common transmembrane helix 6 (TM6) containing interface. Consistent with this, TMD-based FRET studies have revealed sequential agonist-evoked inter- and intra-TMD conformational changes^35,52–54^. Compared to our understanding of agonist-driven activation, the conformational and structural effects of PAMs are less clear. FRET studies have provided strong evidence that PAMs modulate inter-LBD^55^ and inter-TMD conformational changes^29,56^, but a structural framework for interpreting such signals is limited. Most full-length, active state cryo-EM structures incorporate both agonist and PAM for maximal stabilization^30,38,44–46,48^, limiting our ability to disentangle the conformational basis of signaling driven independently by agonist or PAM.

We have recently shown that, in addition to their well-known ability to activate specific heterotrimeric G protein subtypes, a subset of mGluRs, including mGluR3, mGluR7, and mGluR8, is capable of glutamate-evoked desensitization and internalization by GRKs and *β*-arrestins (*β*- arrs)^6,57^. However, the ability of different orthosteric and allosteric compounds to evoke mGluR desensitization has not yet been addressed, raising the question of potential functional selectivity. This knowledge is critical in a therapeutic context as the relative ability of ligands to drive G protein versus GRK/*β*-arr coupling is likely a major determinant of efficacy and long-term compensatory effects, as is well-documented for a variety of family A GPCRs^58–60^.

In this study, we use a combination of live-cell functional assays, measures of conformational dynamics, and single particle cryo-EM to probe the functional and structural basis of allosteric modulation of mGluRs. We find that orthosteric agonist efficacy is correlated with receptor activation and desensitization, while PAMs show a variety of effects which are typically biased towards desensitization. Notably, PAMs can often drive more internalization than glutamate and can even enable glutamate-driven desensitization in mGluR2, a subtype that is otherwise desensitization-resistant. Focusing on mGluR3, FRET analysis reveals a combination of LBD and TMD-level PAM-driven dimeric rearrangements which motivate structural analysis. We thus determine full-length mGluR3 structures via single particle cryo-EM under four distinct ligand conditions, revealing a variety of intra- and inter-domain PAM-driven conformational rearrangements in agonist- and antagonist-bound conditions. Together, our structural analysis reveals multiple inter-TMD interfaces, informing a model of mGluR3 activation and desensitization involving a dynamic population of conformational intermediates differentially stabilized by orthosteric and allosteric ligands.

## Results

### Orthosteric agonists drive activation and desensitization with a correlated maximum efficacy

Motivated by our recent finding that, in addition to initiating G protein signaling,, glutamate binding can drive mGluR desensitization via GRKs and *β*-arrs^57^ we first asked how mGluRs respond to a range of orthosteric agonists (**Fig. S1A**). We initially focused on mGluR3, as this is the subtype for which we previously found the most robust acute desensitization and endocytosis^57^. We assessed the agonists LY379268 (“LY37”) and DCG-IV by first comparing their effects to glutamate in a G protein-dependent G protein-coupled inwardly rectifying potassium (GIRK) current assay (**Fig. 1A**). All three agonists produced dose dependent, reversible GIRK currents (**Fig. 1B; S1B, C**). However, the maximal current evoked by saturating concentrations were larger than glutamate for LY37 and smaller than glutamate for DCG-IV (**Fig. 1C**). These results are consistent with prior studies of group II mGluRs which have characterized LY37 as a “super agonist” and DCG-IV as a partial agonist^39–41,61^.

**Figure 1.**
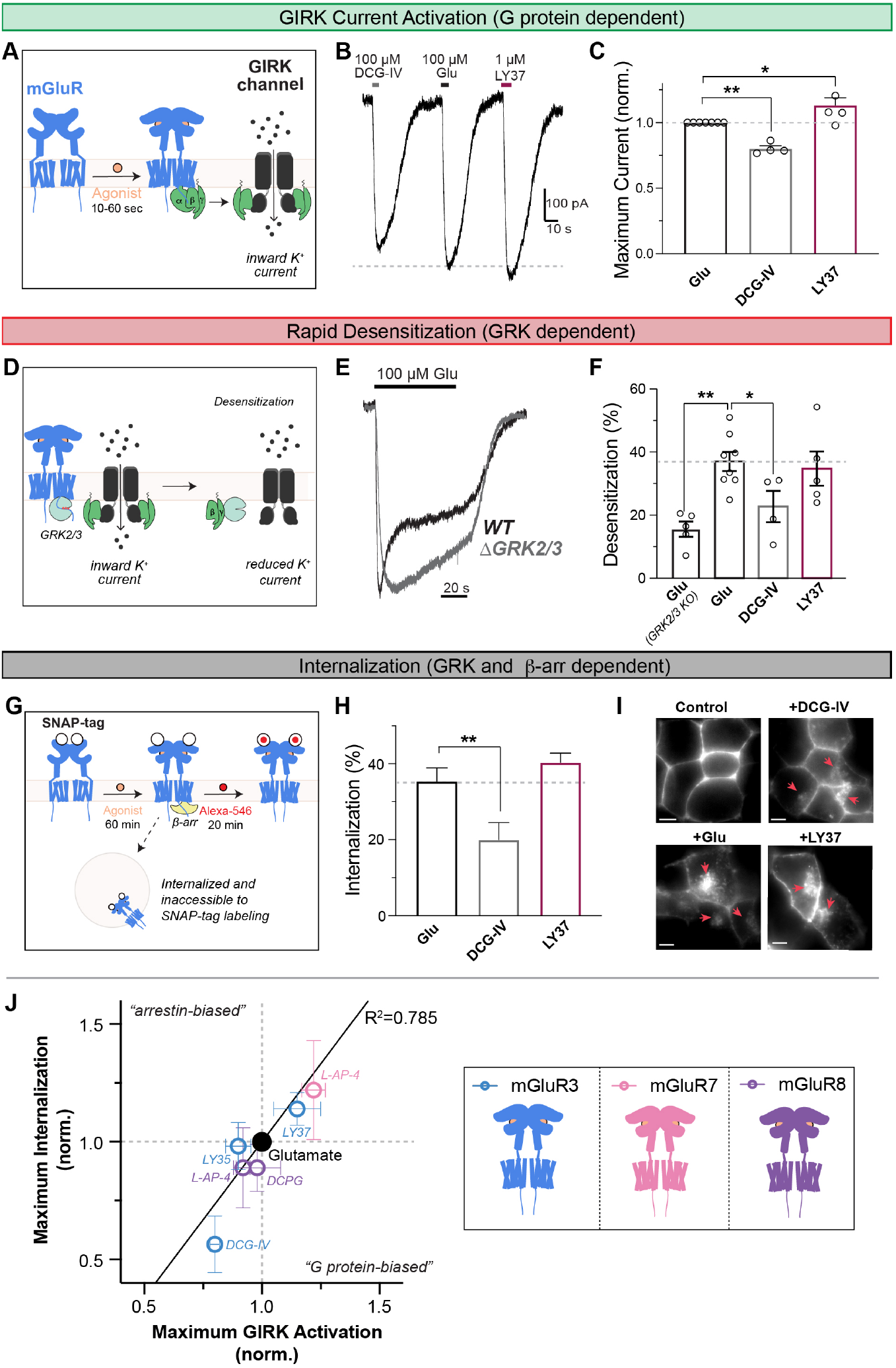
Orthosteric agonists drive activation and desensitization of mGluRs with correlated maximum efficacy. **(A)** Schematic showing mGluR-mediated GIRK activation through Gα_i/o_ activation and Gβγ release. **(B)** Representative whole-cell patch clamp current trace showing agonist-evoked mGluR3-mediated GIRK potassium currents showing the relative maximum agonist efficacies. Grey dashed line represents peak glutamate current. **(C)** Bar plot quantifying the maximum mGluR3-mediated current normalized to the glutamate (100 µM) condition for DCG-IV (100 µM) and LY37 (1 µM). **(D)** Schematic depicting rapid desensitization of mGluR-evoked currents via GRK2/3, which can be recruited via the receptor and, subsequently, scavenge Gβγ to decrease GIRK currents. **(E)** Representative current traces showing acute desensitization of mGluR3-mediated currents in WT HEK293 vs GRK2/3 double KO cells. **(F)** Quantification of desensitization following 60 s application for all the three agonists (same concentrations as panel C). Grey dashed line represents the glutamate mean desensitization value for glutamate. **(G)** Schematic showing surface labelling assay for quantification of ligand-evoked receptor internalization. Cells are incubated with agonist for 60 min followed by labelling of SNAP-tagged surface receptors with membrane impermeable BG-Alexa-546 fluorophore. **(H)** Quantification of the % internalization of mGluR3 following treatment with each agonist (same concentrations as panel C). **(I)** Representative widefield fluorescence images depicting internalization of SNAP-mGluR3 labeled with BG-Alexa-546 prior to agonist treatment for 30 min. Red arrows highlight endocytosed receptors. Scale Bar = 5 µm. **(J)** Scatter plot showing maximum GIRK activation versus internalization for a range of agonists normalized to saturating glutamate for three different mGluR subtypes (mGluR3, mGluR7, mGluR8). Linear regression was applied to the entire data set and constrained to go through the glutamate reference point.

Bar-plots represent mean ± SEM for n>4 cells/condition for electrophysiology experiments and n>30 images per condition (for at least 3 separate days) for imaging experiments.

For C, F, H: One-way ANOVA with multiple comparisons, * p<0.05, ** p<0.01.

See also Figure S1.

We then used the GIRK channel assay to measure acute desensitization of receptor responses following extended (60 s) agonist application (**Fig. 1D**). As previously reported^57^, glutamate drove a peak response that desensitized by ∼40% over the course of 60 s (**Fig. 1E**). This desensitization was markedly reduced in GRK2/3 double knockout (τιGRK2/3) cells (**Fig. 1E, F**). Interestingly, the partial agonist DCG-IV produced weaker desensitization while the super agonist LY37 produced strong, glutamate-like desensitization (**Fig. 1F; Fig. S1D**), suggesting a correlation between agonist efficacy for both activation and desensitization. Consistent with our prior study showing that mGluR2 largely eludes desensitization, glutamate produced negligible desensitization of mGluR2-driven GIRK currents to an identical level in wild type and τιGRK2/3 cells (**Fig. S1E, F**).

To assess agonist-evoked receptor endocytosis, we used our previously established SNAP-tag based surface labeling internalization assay (**Fig. 1G**) where glutamate drives a dose-dependent and β-arr dependent drop in surface fluorescence over the course of 1 hr^57^ (**Fig. S1F, G**). All three agonists drove mGluR3 internalization, but again DCG-IV drove substantially less internalization than glutamate while LY37 drove slightly more (**Fig. 1H**). This same trend was observed upon imaging of cells with fluorophore labeling prior to agonist treatment where a larger accumulation of intracellular receptors was seen following glutamate or LY37 application compared with DCG-IV (**Fig. 1I**). As a control, all three agonists failed to elicit internalization of mGluR2 (**Fig. S1I, J**).

To further test the correlation of agonist efficacy, activation, and desensitization, we extended our analysis to more agonists and receptor subtypes. We tested the group II mGluR agonist LY354740 (“LY35”) and agonists for the group III mGluRs, mGluR7 (L-AP-4) and mGluR8 (L-AP-4, DCPG) in both GIRK activation and internalization assays (**Fig. S1K-P)**. **Fig. 1J** shows a summary of the maximal efficacy of all ligands in both assays, revealing a positive correlation between efficacy of G protein activation and *β*-arr driven internalization. Notably, none of the agonists tested showed a clear bias toward one pathway relative to glutamate as the reference ligand. This suggests that shared agonist-driven conformational changes drive both activation and desensitization and raises the question of how activation and desensitization are related for allosteric ligands.

### PAMs promote activation and desensitization with a bias toward desensitization

PAMs have been reported across mGluR subtypes as potentially viable therapeutics for a range of disorders^25,33,62^, but only one so far has been shown to have activity on mGluR3. This compound, VU6023326 (“VU602”), is a recently reported PAM with similar apparent affinity for mGluR2 and mGluR3^63^. It contains two six membered aromatic rings with bulky substituents connected via a short, ethereal linker, showing general similarity to many mGluR PAMs (**Fig. S2A**). In line with the agonism of mGluR2 we have previously seen with PAMs^29^, VU602 produced dose-dependent GIRK currents upon application to mGluR3-expressing cells (**Fig. 2A, B; Fig. S2B**). The maximal current evoked by VU602 was ∼80% of that produced by glutamate (**Fig. 2B**), consistent with our prior study of mGluR2, where all PAMs tested showed a 60-80% efficacy^29^. We then asked if application of PAM alters the maximal response to agonists by applying VU602 in the presence of saturating glutamate or LY37. In both cases, VU602 produced a clear inward current that boosted the maximum response to glutamate or LY37 by 20-30% (**Fig. 2C, D; Fig S2C, D**). This result reveals that mGluR PAMs can not only serve as allosteric agonists and modulators of agonist affinity but can also boost agonist efficacy.

**Figure 2.**
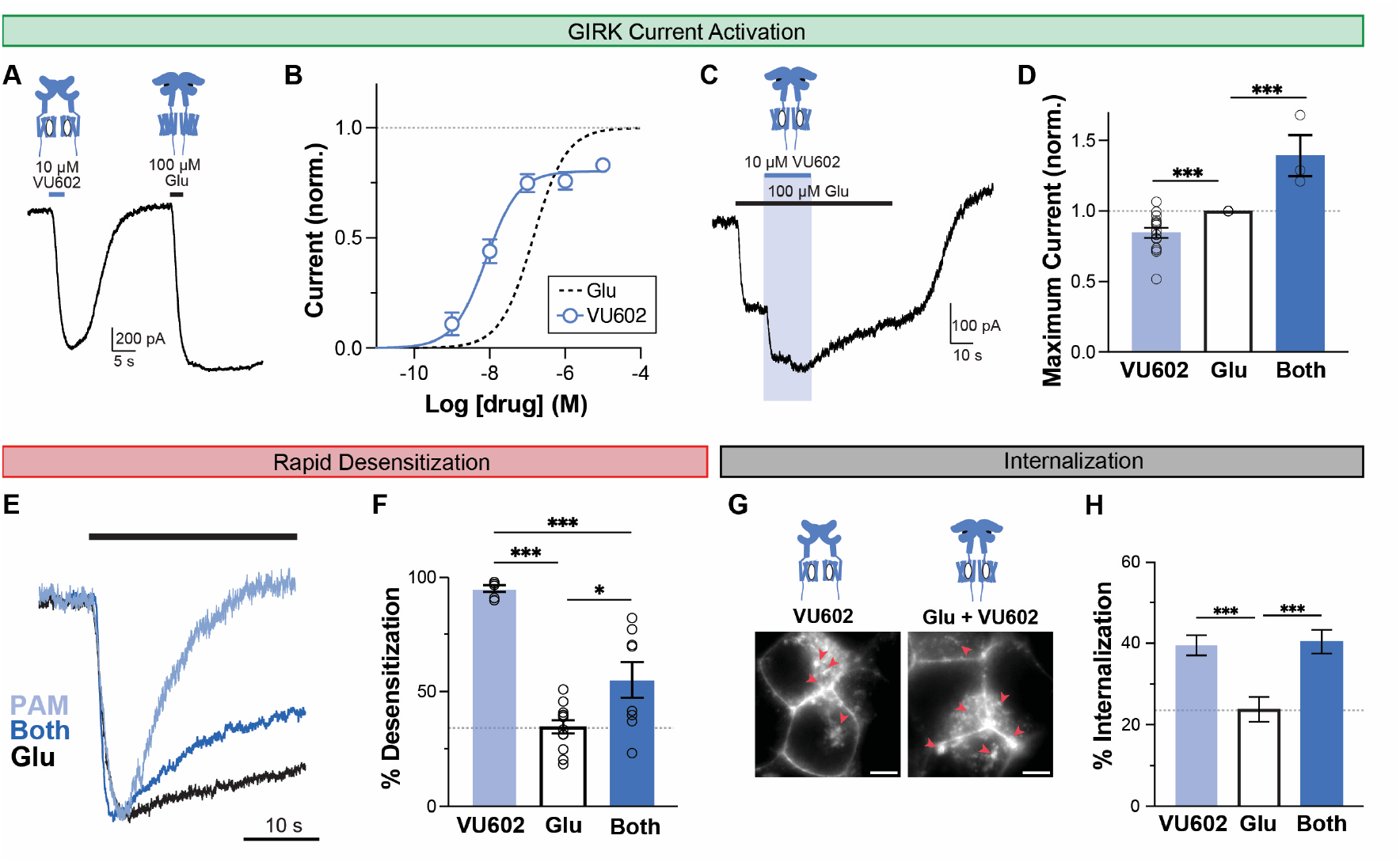
An mGluR3 PAM serves as a partial agonist, boosts the maximal response to glutamate, and strongly drives desensitization. **(A)** Representative GIRK current trace showing the response to a saturating dose of VU602 (10 µM) compared to glutamate. **(B)** Dose response curve for VU602-evoked GIRK currents normalized to the response to saturating glutamate. EC_50_ = 10.8 ± 4.6 nM. Black dashed line shows the glutamate dose response curve as a reference. **(C)** Representative GIRK current trace showing that application of VU602 in the presence of saturating glutamate further potentiates mGluR3-driven currents. **(D)** Bar plot summarizing the maximum response for VU602 (10 µM), glutamate (100 µM), or co-application of both. Glutamate is used as the reference for normalization. **(E)** Representative traces showing acute GIRK current desensitization following application of VU602 (10 µM), glutamate (100 µM), or both. **(F)** Quantification of the extent of desensitization for the three conditions in E. **(G)** Bar plot showing mGluR3 internalization in response to VU602 (10 µM), glutamate (100 µM), or co-application of both. **(H)** Representative images showing VU602 and VU602+Glu evoked internalization. Scale bar = 5 µm. Bar-plots represent mean ± SEM for n>3 cells for electrophysiology experiments and n>40 images per condition (for at least 4 separate days) for imaging experiments. For D, F, H: One-way ANOVA with multiple comparisons, *** p<0.001 See also Figure S2.

We then characterized the effects of VU602 on mGluR3 desensitization. Strikingly, acute desensitization of mGluR3-driven GIRK currents was enhanced for currents evoked either by VU602 alone or VU602 and glutamate together, compared to glutamate alone (**Fig. 2E, F; Fig. S2E**). In addition, VU602 produced a larger extent of mGluR3 internalization than glutamate and the response was slightly boosted when ligands were co-applied (**Fig. 2G, H; Fig. S2F**). Importantly, VU602-driven mGluR3 internalization was *β*-arr dependent (**Fig. S2G**). Together these data indicate that VU602 is capable of both initiating G protein-dependent activation and GRK and *β*-arr-dependent desensitization, but that the relative efficacy is higher for desensitization.

Motivated by the apparent desensitization bias of VU602 on mGluR3, we tested a panel of PAMs on group II and III mGluRs in both GIRK activation and internalization assays (**Fig. 3**). First, we tested the group III PAM VU6005649 (“VU600”)^64^. In the GIRK activation assay, VU600 served as a partial agonist for mGluR7 with ∼80% efficacy versus glutamate and also boosted the response to saturating 10 mM glutamate (**Fig. 3A, B; Fig. S3A, B**). In the surface labeling assay, VU600 produced almost twice as much internalization as 10 mM glutamate and co-application led to an even larger drop in surface fluorescence (**Fig. 3C**). The enhanced internalization produced by VU600 compared to glutamate could be clearly visualized in live cells (**Fig. 3D**). When investigating mGluR8, VU600 showed a similar profile in both assays but with a slightly less pronounced difference compared to glutamate in internalization assays (**Fig. 3E-H; Fig. S3C, D**). We also tested the mGluR8 PAM AZ 12216052 (“AZ122”)^65^ which similarly showed partial agonism in the GIRK current assay, although it did not produce a substantial boost of the response to saturating glutamate (**Fig. 3F**). In internalization assays, AZ122 showed similar results to VU600 with stronger internalization than saturating glutamate and a boost upon drug co-application (**Fig. 3G, H**).

**Figure 3.**
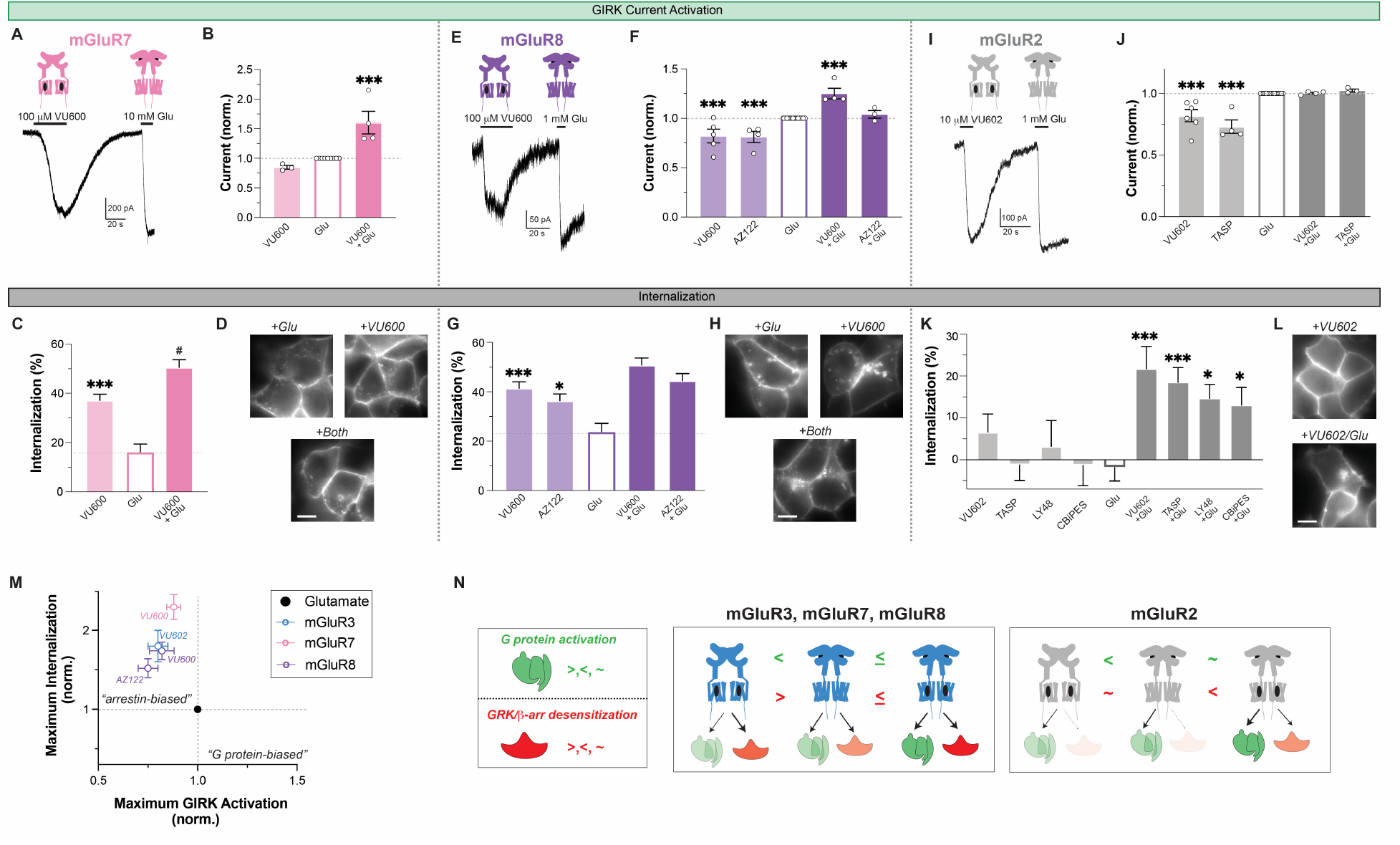
PAMs show desensitization bias across the group II/III mGluR subfamilies. **(A)** Representative trace showing currents elicited by saturating VU600 PAM (100 µM) and glutamate (10 mM) in cells expressing mGluR7. **(B)** Bar plot showing normalized maximum mGluR7-mediated current for VU600 (100 µM), glutamate (10 mM) or the combination of both. **(C-D)** Bar plot (C) and representative images (D) showing extent of mGluR7 internalization for VU600 (65 µM), glutamate (10 mM), or both. **(E)** Representative trace of elicited currents by saturating VU600 (100 µM) vs saturating glutamate (1 mM) for mGluR8. **(F)** Bar plot showing normalized maximum mGluR8-mediated current for VU600 (100 µM), AZ122 (100 µM), glutamate (1 mM) or the combination of either PAM with glutamate. **(G-H)** Bar plot (G) and representative images (H) showing extent of mGluR8 internalization for VU600 (260 µM), AZ122 (100), glutamate (1 mM), or the combination of either PAM with glutamate. **(I)** Representative trace showing currents elicited by saturating VU602 PAM (10 µM) vs saturating glutamate (1 mM) for mGluR2. **(J)** Bar plot showing normalized maximum current for VU602 (10 µM), TASP (10 µM), glutamate (1 mM), or the combination of either PAM with glutamate. **(K-L)** Bar plot (K) and representative images (L) showing extent of mGluR2 internalization for saturating doses of mGluR2 PAMs VU602 (10 µM), TASP (10 µM), LY48 (10 µM) and CBiPES (10 µM), glutamate (1 mM) or PAMs + glutamate. Note: GRK2 was co-expressed for mGluR2 internalization experiments. **(M)** Summary plots of normalized maximum GIRK activation versus maximum internalization for all PAMs used in this study on mGluR3, mGluR7, or mGluR8 with all values normalized to the effects of saturating glutamate. **(N)** Schematics summarizing the effects or PAMs, glutamate or both on G protein activation versus GRK/*β*-arr desensitization across different mGluR subtypes. Bar-plots represent mean ± SEM for n>3 cells for electrophysiology experiments and n>30 images per condition (>3 days) for imaging experiments. For panels B, C, F, G, J, K, One-way ANOVA with multiple comparisons; * p<0.05, *** p<0.001. See also Figure S3

We next asked what the effects of PAMs are on mGluR2, a subtype which is resistant to agonist-driven desensitization (**Fig. S1H, I**)^57^. We found that VU602 produced clear dose-dependent partial agonism of mGluR2 in the GIRK current assay, consistent with what we have seen for a panel of mGluR2 PAMs^29^ (**Fig. 3I, J; Fig. S3E**). VU602 showed a similar efficacy to TASP0433 (“TASP”), the mGluR2 PAM which showed the highest efficacy in our prior study. Interestingly, both VU602 and TASP failed to further enhance the saturating glutamate response (**Fig. 3J; Fig. S3F**). In the internalization assays, neither PAM nor glutamate evoked any clear effects (**Fig. 3K**). However, co-application of PAM and glutamate produced a clear internalization of mGluR2 with the largest effects for VU602 and TASP (**Fig. 3K, L**). Importantly, the Glu/PAM-evoked internalization of mGluR2 was GRK and *β*-arr-dependent (**Fig. S3G**), revealing that PAMs can drive *β*-arr coupling even in subtypes that do not typically recruit *β*-arr in response to agonists. We also tested the ability of PAMs to drive acute desensitization of mGluR2-evoked GIRK currents and found that co-application of TASP and glutamate produced a substantial increase in the speed and extent of desensitization in a GRK2-dependent manner (**Fig. S3H-J**).

Together, these data show that, while there is variability across mGluR PAMs, they tend to show a common profile. All PAMs serve as partial agonists toward G protein with a subset that can substantially boost the maximal efficacy of glutamate. In terms of desensitization, all PAMs tested showed a bias toward desensitization with most driving stronger internalization than glutamate alone. **Fig. 3M** shows a summary of the relative efficacy of G protein activation (i.e. maximal GIRK current amplitude) versus *β*-arr coupling (i.e. maximum internalization) for all mGluR3, mGluR7, and mGluR8 PAMs tested, highlighting a clear *β*-arr bias. **Fig. 3N** summarizes the major effects of PAMs versus glutamate versus both on relative G protein and *β*-arr coupling.

### PAMs drive LBD-dependent and -independent conformational and functional effects

To probe the conformational effects of allosteric modulation, we turned to established inter-subunit mGluR FRET sensors that detect activation-associated rearrangement between subunits either at the LBDs^39^ or TMDs^56^. We previously found that PAM application alone had no effect on inter-LBD FRET in mGluR2, but that it could enhance the response to agonists^29^. In contrast to this result, application of VU602 produced a modest, but clear decrease in inter-LBD FRET when applied to SNAP-mGluR3 expressing cells (**Fig. 4A, B; Fig. S4A, B**). VU602 did not produce a response in SNAP-mGluR2 expressing cells (**Fig. S4C**). We reasoned that this subtype discrepancy may be due to the elevated basal inter-LBD conformational dynamics of mGluR3 which mimic the agonist-bound state^41,66^. Consistent with this, introduction of the S152D mutant, which blunts basal inter-LBD dynamics^41^, or application of the orthosteric antagonist LY341495 (“LY34”) reduced and abolished the VU602 response, respectively (**Fig. S4C**; **Fig. 4C**). The NAM MNI137 also fully blocked the VU602 response (**Fig. 4C**), likely through a competitive binding mechanism. When co-applied with glutamate, VU602 did not further boost the maximal response (**Fig. 4B**) as it did in the GIRK current assay (**Fig. 2C**).

**Figure 4.**
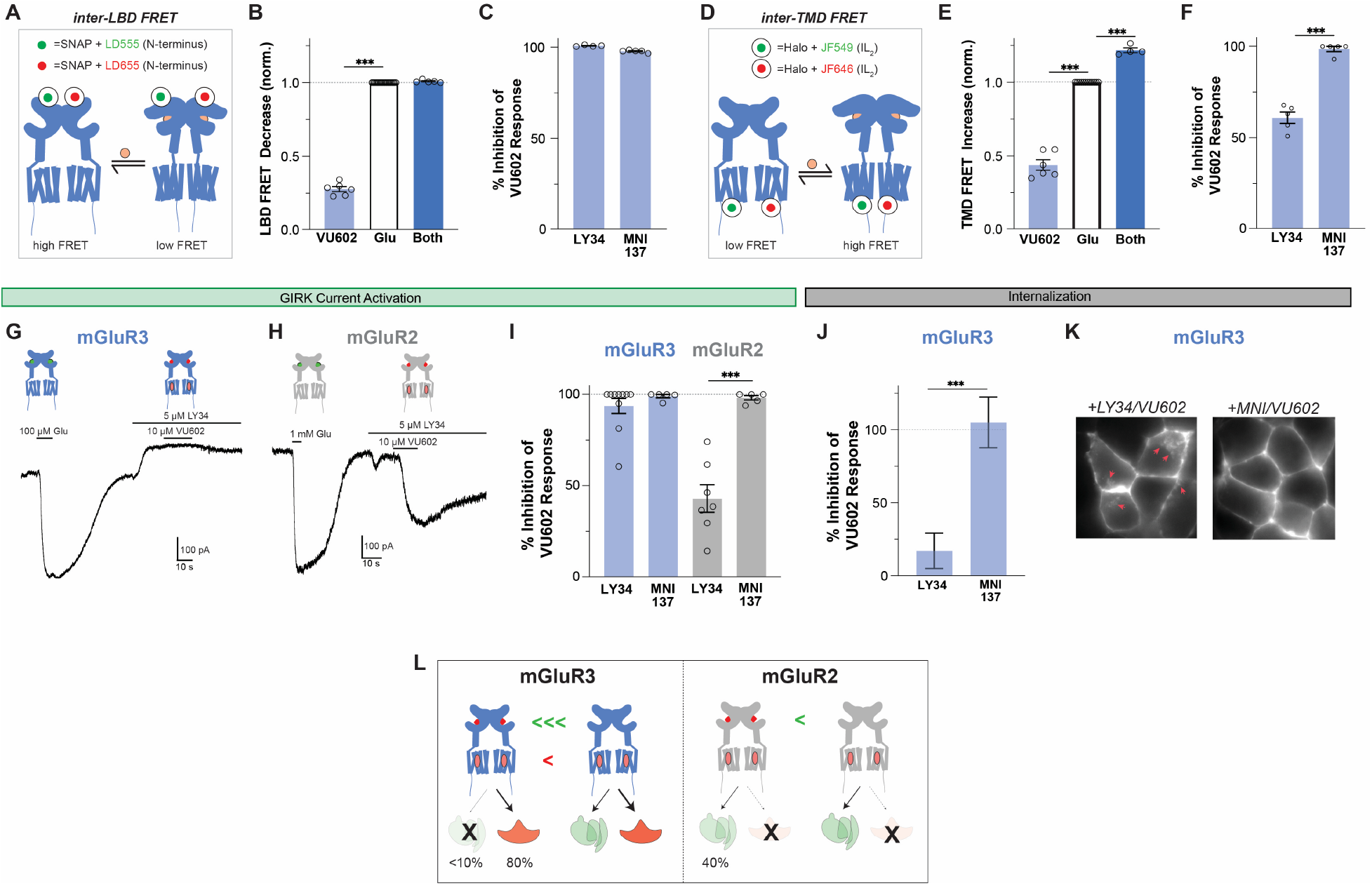
PAMs produce LBD-dependent and independent conformational and functional effects. **(A)** Schematic showing inter-LBD FRET sensor where LBD closure and reorientation from “relaxed” to “active” dimer states upon agonist exposure decrease FRET efficiency. **(B)** Bar graph showing normalized mGluR3 inter-LBD FRET decrease following application of saturating VU602 (10 µM), glutamate or both. **(C)** Bar graph showing the inhibition of VU602 inter-LBD FRET responses by competitive antagonist (5 µM LY34) or NAM (60 µM MNI-137). **(D)** Schematic showing inter-TMD FRET sensor where agonist application receptor increases the FRET efficiency as TMD domains undergo activation-associated rearrangements. **(E)** Bar graph showing normalized mGluR3 inter-TMD FRET increase following application of saturating VU602 (10 µM), glutamate or both. **(F)** Bar graph showing the inhibition of VU602 inter-TMD FRET responses by saturating competitive antagonist (10 µM LY34) or NAM (60 µM MNI-137). **(G)** Representative GIRK current trace for mGluR3 showing the lack of response to VU602 when applied in the presence of competitive antagonist LY34. **(H)** Same as in G but for mGluR2, which maintains a clear VU602 response in the presence of LY34. **(I)** Bar graph summarizing VU602 evoked GIRK currents when applied in the presence of the antagonist LY34 (5 µM) or NAM MNI137 (10 µM) for mGluR3 and mGluR2. Values are normalized to the response to saturating glutamate. Dotted line shows the response to VU602 alone for each receptor. **(J)** Bar graph showing VU602-induced internalization for mGluR3 in the presence of the antagonist LY34 (5 µM) or NAM MNI137 (10 µM). Dotted line shows the response to VU602 alone. **(K)** Representative images showing mGluR3 internalization in the presence of antagonist and PAM but not NAM and PAM conditions. Red arrows highlight clusters of internalized receptors. **(L)** Schematic summarizing the effects of a competitive antagonist on PAM-evoked activation and desensitization of mGluR3 and mGluR2. Bar-plots represent mean ± SEM for n> 5 separate preparations for FRET experiments, n>5 cells for electrophysiology experiments, n>30 images per condition (for at least 3 separate days) for imaging experiments. For **I** t-test, *** p<0.001; n.s. p≥0.05. For **B** and **E**, One-way ANOVA with multiple comparisons; For **F** and J, unpaired T-test. ** p<0.01, *** p<0.001. See also Figure S4.

We next turned to inter-TMD FRET (**Fig. 4D**) where we found that VU602 can elicit a clear, partial response when applied alone (**Fig. 4E; Fig. S4D, E**). Notably, the relative inter-TMD PAM FRET response was larger (∼45% versus saturating glutamate) compared to the inter-LBD PAM FRET response (∼30% versus saturating glutamate). The inter-TMD FRET response was only partially reduced by saturating LY34 application but was nearly abolished by MNI137 (**Fig. 4F**). Unlike what was seen with inter-LBD FRET, VU602 was also able to boost the response to saturating glutamate (**Fig. 4E**). Together these data show that PAMs can drive inter-LBD and inter-TMD conformational changes. However, only for the TMD sensor, do PAMs boost the maximal response to agonist or report a conformational change in the presence of a competitive antagonist suggesting a larger effect on TMD than LBD conformation.

Given the ability of PAMs to drive conformational changes that are independent of LBD closure and rearrangement, we asked what the functional properties are of antagonist/PAM-bound states. Strikingly, we found that pre-application of LY34 abolished the GIRK current response to VU602 in mGluR3-expressing cells (**Fig. 4G**). This contrasts with mGluR2, which was able to drive a partial GIRK response to VU602 in the presence of saturating LY34 (**Fig. 4H**), consistent with our prior study of mGluR2 PAM-driven currents^29^. Both mGluR2- or mGluR3- mediated VU602 responses were fully blocked by the NAM MNI137 (**Fig. S4F, G**). These results suggest that distinct conformational coupling properties between TMDs and LBDs exist in different mGluR subtypes.

We then asked how antagonist application would alter PAM-driven mGluR internalization. Unlike what was seen with GIRK currents, LY34 was only able to weakly inhibit VU602-driven internalization of mGluR3 (**Fig. 4J, K; Fig. S4H**). In key controls, LY34 fully blocked glutamate-driven internalization (**Fig. 4J**) and MNI137 fully blocked both VU602 and glutamate-driven internalization (**Fig. S4H, I**). The ability of a PAM to drive mGluR3 internalization, but not G protein-activation, in the LY34-bound state indicates that a GRK/*β*-arr biased conformation may be favored. Overall, these studies show both ligand and subtype-specific effects, suggesting underappreciated conformational complexity in mGluR activation and desensitization (**Fig. 4L**).

### Agonist-bound cryo-EM structures of mGluR3 with and without PAM reveal PAM-evoked structural reshaping at key interfaces

Motivated by the complex functional and conformational effects of PAMs, we turned to single particle cryo-EM to gain more insight into the activation and desensitization-associated structural states of mGluRs. We aimed to solve structures of mGluR3 with and without VU602 to enable the first comparative analysis of an mGluR structure with and without PAM. To avoid complications due to the conformational dynamics seen at the LBD level in un-liganded mGluR3^41,66^, we prepared four separate samples with either antagonist (LY34) or agonist (LY37) alone or in combination with VU602.

We first determined the structure of detergent-solubilized, full length mGluR3 in the presence of LY37 alone or both LY37 and VU602 (**Fig. 5A**). To facilitate direct comparison, the same purification and grid-freezing protocol was used for all samples. Initial processing resulted in full-length structures with a global resolution of 3.5 Å and 3.3 Å for LY37 and LY37/VU602 conditions, respectively (**Fig. S5**). Consistent with prior full-length mGluR structures^30,38,44–48^, we obtained higher resolution at the LBDs with resolution decreasing moving down the receptor to the CRDs and TMDs with no density observed for CTDs. We enforced C2 symmetry at the last stage of processing the full-length map (**Fig. S5**), as it improved resolution while maintaining all features seen without symmetry constraints. As we anticipated that flexibility between the extracellular domain (LBD and CRD) and the TMDs contributes to heterogeneity within our full-length map, we performed local refinement for the extracellular domain of the receptor. At the TMD level, we first performed particle subtraction to isolate the TMDs followed by non-uniform refinement, maintaining C2 symmetry. This approach did not improve reported TMD resolution but did reveal additional density for TM helices that were not well resolved in the full-length map (**Fig. S5**). LY37 only and LY37/VU602 structures share a similar global domain arrangement to each other, and to other agonist/PAM bound mGluR structures^30,38,44,46,47^, with increasing RMSD from LBDs to CRDs to TMDs (**Fig. 5A-C; Fig. S6A).** Due to the high resolution at the LBDs, we were able to clearly place the LY37 agonist in both structures and to assign density to ions (**Fig. S6B, C**). These ion densities appear in overlapping locations to Cl^-^ ions seen in an mGluR3 LBD crystal structure^67^, although we cannot unambiguously define their identity.

**Figure 5.**
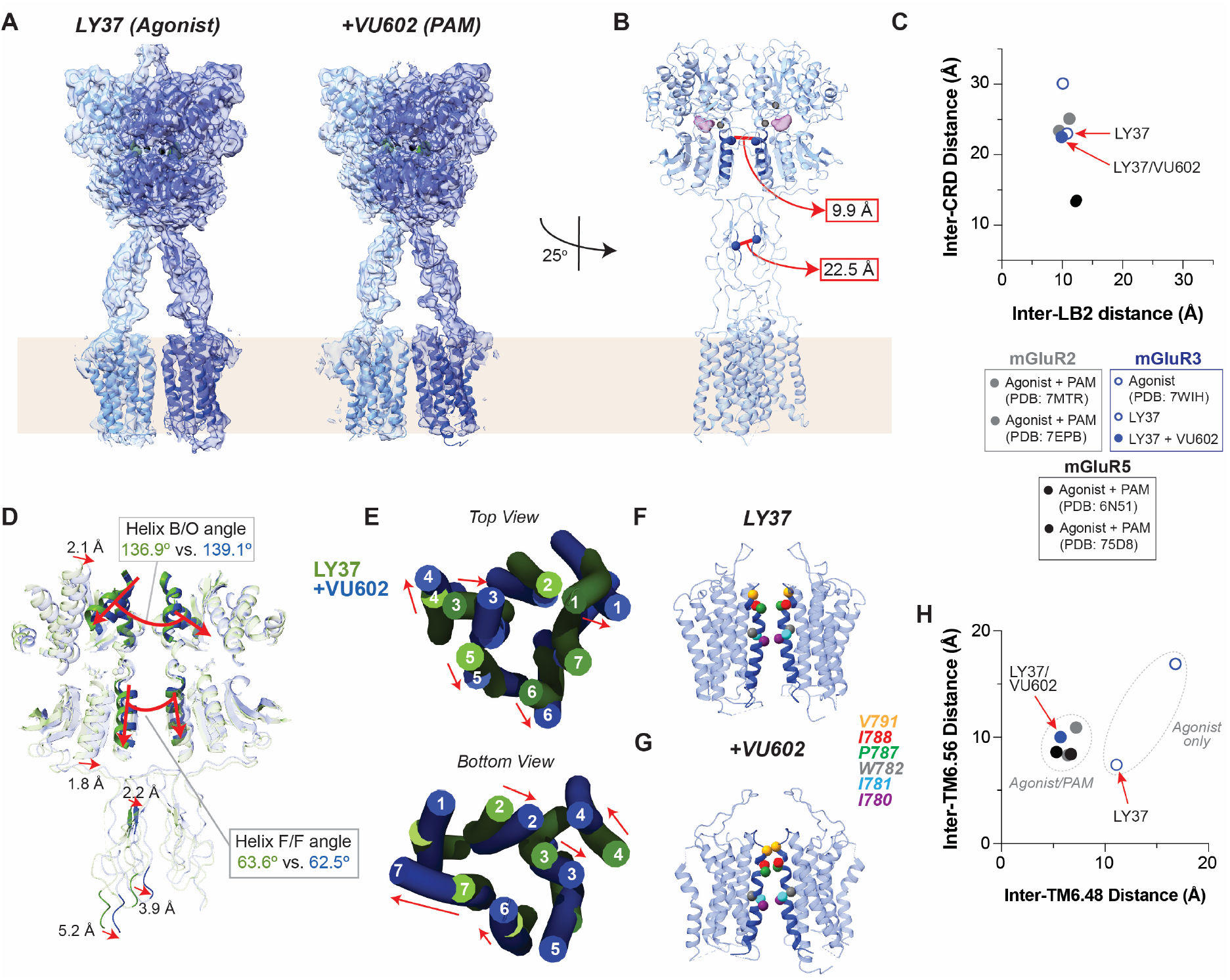
Agonist-bound structures of mGluR3 show common intersubunit interfaces that are re-shaped by allosteric modulators. **(A)** Full length cryo-EM density map and protein model for LY37 (agonist) and LY37/VU602 (agonist/PAM) bound mGluR3. **(B)** Full-length mGluR3 LY37/VU602 model with LY37 in purple and ions in black highlighting the inter LBD bottom lobe (LB2) distance measured from the Cα of E224 and inter-CRD distance measured from the Cα of C546. **(C)** Plot comparing inter-LB2 versus inter-CRD distance across reported agonist-bound homodimeric mGluR subtypes measured via conserved positions highlighted in (B). **(D)** mGluR3 LY37 (green) and mGluR3 LY37/VU602 (blue) structures aligned via the top lobe (LB1) of chain A. The angle between helix B and helix O and the angle between helix F of chain A and chain B is reported. Structural offsets are reported for residues T410 (2.7 Å), R206(1.8 Å), I530(2.2 Å), M547(3.9 Å), and E567 (5.2 Å). **(E)** Top and bottom views of single TMD alignments between LY37 and LY37/VU602 structures with offsets highlighted by red arrows. Alignments are performed for the entire TMD from residue 571 to the end of the model, including loops. **(F-G)** Side views of LY37 (F) and LY37/VU602 (G) TMD dimer with TM6 shown in dark blue and the Cα of key residues highlighted. **(H)** Plot of inter TM6 distance across full-length homodimeric mGluR structures using two conserved positions. See also Figures S5 and S6

In both structures, the LBDs are stabilized in a closed/closed-active conformation with a similar inter-LB2 distance to prior agonist-bound cryo-EM structures of mGluRs (**Fig. 5B, C**). Furthermore, both structures show compact CRDs with an inter-subunit distance of ∼20-25 Å (**Fig. 5B, C**). An analysis of inter-CRD distance across agonist-bound mGluR structures reveals more variability compared to inter-LBD distances, with our mGluR3 structures more closely resembling mGluR2 than mGluR5 structures (**Fig. 5C**). A closer analysis of the LBDs revealed subtle rearrangements in the LY37/VU602-bound structure compared to LY37 only, which may explain the effects of PAM on agonist affinity and efficacy (**Fig. 5D**). In the presence of VU602, a compaction occurs at the LBD dimer interface, characterized in the upper lobe (LB1) as a change in angle between helix B of one protomer and helix O of the other protomer, from 136.9° to 139.1° in the LY37 and LY37/PAM structures, respectively (**Fig. 5D**). At the lower lobe (LB2), a change in angle from 63.6° (LY37) to 62.5° (LY37/VU602) is seen between helix F on each protomer, ultimately creating a tighter inter-LBD interface (**Fig. 5C, D**). Most importantly, this inter-LBD rearrangement reshapes the LY37 binding site. While the same LBD residues interact with LY37 in both structures, the top lobe residues S151 and R68 are closer to the ligand in the LY37/VU602 structure (**Fig. S6B**). This change in ligand binding cleft compaction can also be measured as a decrease in inter-lobe distance in the LY37/PAM condition between LB2 residue Y222 and LB1 residues A172, S173, and T174 (**Fig. S6B**). Overall, our structures show that subtle changes in LBD compaction and cleft closure occur with the addition of VU602.

Differences are also observed in CRD compaction between the LY37 and LY37/VU602 structures, with a general CRD rearrangement that decreases the inter-subunit distance by 2-5 Å in the LY37/VU602 structure (**Fig 5D**). Finally, extracellular loop 2 (EL2), also exhibits a reshaping and positional shift in the presence of PAM (**Fig. S6D, E**), which may contribute to inter-domain allostery as has been proposed in previous mGluR structures^38,44,46^.

While we do not have the resolution to identify VU602 density, the VU602-containing structure shows intra- and inter-subunit rearrangement of the TMD. This includes a widening of the extracellular opening of the TMD bundle due to an outward shift of TM1, TM5, and TM6, and a repositioning of TM3, in the VU602-containing structure (**Fig. 5E; Fig. S6F**). A re-organization is also seen at the intracellular face of the TMD, with offsets for TM1, TM2, TM3, TM4, and TM6, which moves inward and away from the interface in the VU602-containing structure (**Fig. 5E; Fig. S6F**). Furthermore, in the LY37/VU602 model TM7 shows lengthened helical character of about one turn on the extracellular end and an outward bend on the intracellular end compared to the LY37 model (**Fig. S6F**). Together these conformational changes likely reposition intracellular loops, a key step in transducer coupling (**Fig. 5E**)^45,46,48^. An existing mGluR2 structure has resolved PAM density in the absence of G protein (PDB: 7MTR), where the PAM ADX55164 is clearly seen in one TMD^46^. When aligning our mGluR3 TMD structures with the mGluR2 PAM-containing TMD, we observe that the TM helix shifts observed in our LY37/VU602 structure enable a closer alignment with the PAM-bound mGluR2 TMD than seen with our LY37 only structure (**Fig. S6G**). This includes opening of an apparent cavity on the extracellular face of the TMD between TM3, TM5, and TM6 where ADX55164 is observed. Using a cavity detection tool on complete receptor models (see *Methods* section for details), it is apparent that VU602 can fit between TM5 and TM6 in the LY37/VU602 structure, but no cavity is detected in the same region in the LY37 only structure, supporting the conclusion that VU602 binds and causes a conformational rearrangement within the TMD (**Fig. S6H**).

A TM6 containing dimer interface is observed in both structures (**Fig. 5F, G**), consistent with agonist/PAM-bound structures from other mGluR subtypes^30,38,44,46,47^ and the high degree of TM6 conservation across mGluRs (**Fig. S6I**). However, between mGluR3 structures we observe a difference in both the shape of TM6 and the inter-TMD interface profile. In the absence of VU602, TM6 curves such that its closest inter-TMD distance occurs at the middle of TM6 and then widens towards the extracellular face. In contrast, the VU602-containing structure shows a straighter TM6 with the tightest inter-TMD distance occurring at the most extracellular region of TM6, as seen in other agonist/PAM mGluR structures (**Fig. 5H; Fig. S6J**). This suggests that PAM-containing TMDs have a distinct interface compared to agonist only bound receptors. This finding is consistent with inter-TMD measurements and TM6 shape from other ligand-stabilized family C GPCR structures (**Fig S6K, L**). Notably, a previously reported agonist-bound mGluR3 structure^47^ observed a particularly large inter-TM6 distance that diverges with what is seen in ours and other full-length mGluR structures (**Fig. 5H**).

Reshaping of the TMD by PAMs is also supported by three dimensional variability analysis (3DVA), a powerful tool previously used in the analysis of GPCRs^68–72^ and other dynamic proteins^73^. We performed this analysis for both LY37 and LY37/VU602 final particle sets. Overall, 3DVA identified more distinct motions for the LY37 structure compared to the LY37/VU602 structure, suggesting that the LY37/VU602-bound state is less dynamic. Motions are largely observed at the TMDs and CRDs, while the LBDs remain relatively stable in both states (**Fig. S6M**). One motion component only observed from the LY37 data set shows a straightening and compaction of the top of TM6 and the CRDs (**Fig. S6L; supplemental Video 1**), suggesting that LY37 bound mGluR3 may sample different TM6-containing interfaces and the addition of VU602 stabilizes one of the more compact states. Additionally, the LY37 data reveals highly dynamic TMD density in which breathing at the TMD dimer interface is observed such that TM5 moves towards and away from the interface (**Fig. S6L; supplemental Video 2**). This motion may be suggestive of a rolling interface from TM5 to TM6 along the activation pathway. Both LY37 and LY37/VU602 data sets show movement of CRDs relative to each other such that they rotate from being directly across from each other to one passing the other, which is consistent with the two interconverting CRD FRET states observed for mGluR2 in the presence of saturating agonist^49^. (**Fig. S6M; supplemental Video 3 and 4)**. Overall, this analysis suggests that the LY37 data set samples a range of conformations and the addition of VU602 stabilizes a subset of these states.

### Antagonist-bound cryo-EM structures of mGluR3 with and without PAM reveal multiple CRD and TMD positions accommodated by open LBDs

To gain structural insight into the effects of PAMs in the absence of agonist, we next prepared samples with the antagonist LY34 with or without VU602. We reasoned that a VU602 only condition would be difficult to interpret and likely highly dynamic (i.e. low resolution), due to the basal dynamics of the mGluR3 LBD^41,66^. In addition, we anticipated that the LY34/VU602 ligand combination may provide an opportunity to stabilize intermediates along the activation/desensitization pathway.

Following protein purification and grid preparation with the same protocols used for LY37-bound structures, we obtained structures of detergent-solubilized, full length, mGluR3 with LY34 in the absence and presence of VU602 (**Fig. 6A; Fig. S7**). As with LY37-bound structures, we first obtained a high-resolution full-length map and then performed local refinement at the ECD and particle subtraction followed by non-uniform refinement at the TMDs (**Fig. S7**). The LY34 only condition resulted in a heterogeneous data set with three distinct full length cryo-EM density maps (**Fig. 6A**). All three classes show open LBDs in a relaxed inter-subunit conformation, as expected given the presence of an orthosteric antagonist. The “class 1” map represents a relatively high resolution (global resolution=3.4 Å) asymmetric state with a TM3 and TM4-containing interface. The “class 2” map contains high resolution LBDs (local resolution=3.4 Å) but poor resolution at the CRDs and TMDs. The best resolved state (global resolution= 3.2 Å), “class 3”, is asymmetric and shows a TM5-containing TMD interface.

**Figure 6.**
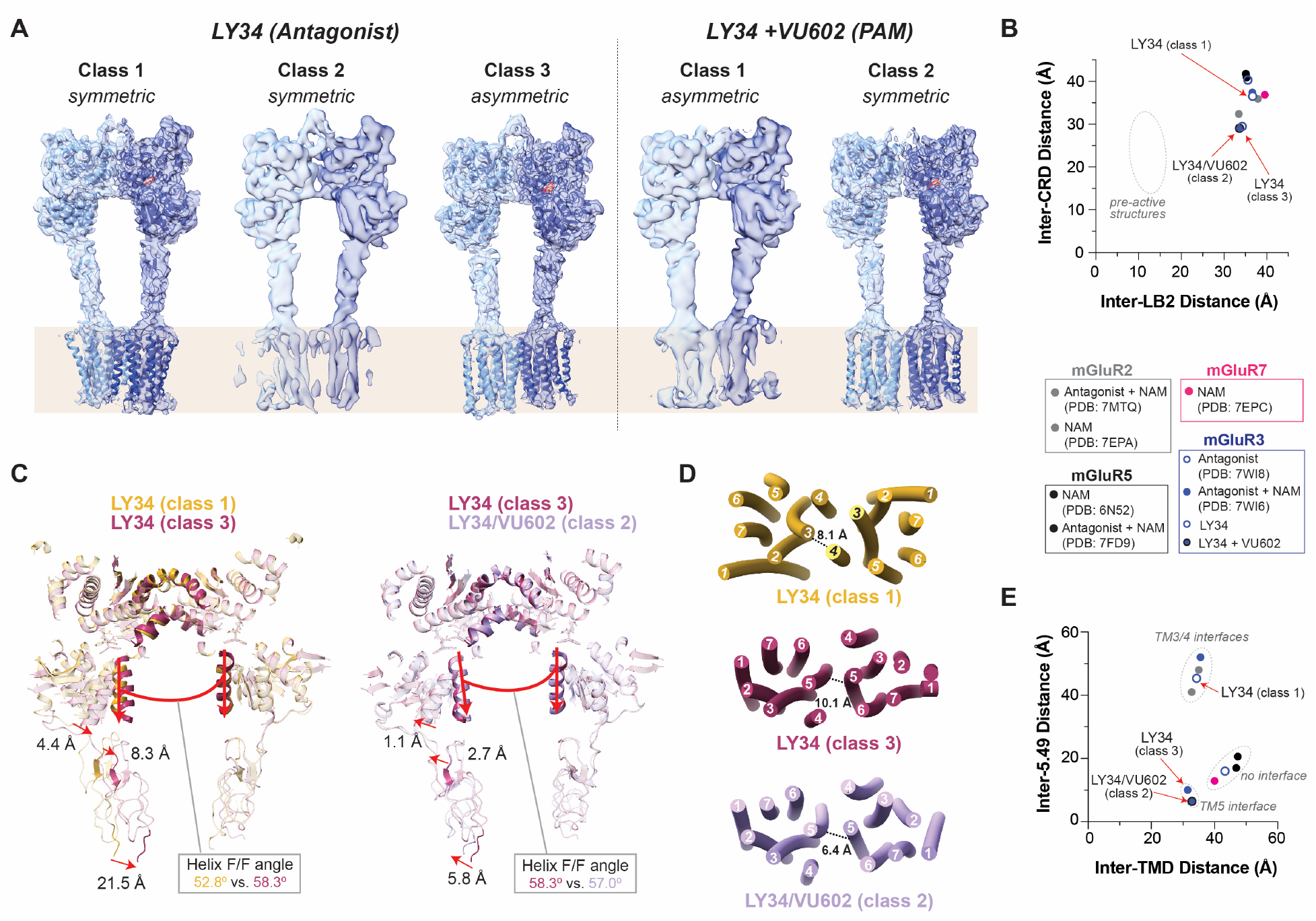
Antagonist-bound structures of mGluR3 reveal a dynamic inter-CRD positioning and multiple TM3/4/5 inter-TMD interfaces. **(A)** Full-length cryo-EM density maps and protein model, when applicable, for LY34 (antagonist) and LY34/VU602 (antagonist/PAM) bound mGluR3. LY34 class 2 and LY34/VU602 class 1 density maps are depicted with a low pass filter of 6 Å. **(B)** Plot comparing inter-LB2 versus inter-CRD distance across reported antagonist and or NAM-bound homodimeric mGluR subtypes measured via conserved positions as in Fig. 5B. **(C)** LY34 class 1 (gold), LY34 class 3 (raspberry) and LY34/VU602 class 2 (lavender) structures aligned via the top lobe (LB1) of chain A. Structural offsets are reported for residues, R206(4.4 Å and 1.1 Å), I530 (8.3 Å and 2.7 Å), and P566 (21.5 Å and 5.8 Å). **(D)** Top view of TMDs with inter-TMD distance measured from the alpha carbon of Cα V639 (TM3) and I708 (TM4) for LY34 class 1 and between V746 (TM5) for LY34 class 3 and LY34/VU602 class 2. **(E)** Plot of inter-TM5 distance across full length homodimeric mGluR structures using a conserved TM5 position (y-axis) and the center of mass distance of the TMD bundles (x-axis). See also Figures S7 and S8

The LY34/VU602 data set also represents a heterogeneous, but more structurally similar, ensemble of states. We resolved two distinct full length cryo-EM density maps with equivalent particle numbers. “class 1” is a lower resolution asymmetric map with somewhat poor TMD density, while the “class 2” map is higher resolution (global resolution= 3.3 Å) and C2 symmetric (**Fig. 6A**). Both class 1 and class 2 maps appear to show TM5-containing interfaces, although the improved resolution makes this unambiguous only for class 2.

The high-resolution class 1 and class 3 LY34 structures show major differences in domain orientations, leading to a global RMSD of 7.40 Å, with much of the difference stemming from the TMD position (**Fig. S8A**). In contrast, the LY34 class 3 structure produces a much better global alignment with the LY34/VU602 class 2 structure with a global RMSD of only 1.36 Å (**Fig. S8A**). LY34-bound mGluR3 structures show similar LBD conformations to each other and to previously reported antagonist and/or NAM-bound mGluR structures, as measured by LB2 distance (**Fig. 6B**). Similarly, all antagonist and/or NAM bound structures show larger inter-CRD distances than observed for agonist bound structures (**Fig. 6B**), but there is more variability in this measurement across structures with the closest inter-CRD distance observed for LY34 class 3 and LY34/VU602 class 2 (**Fig. 6B**). Alignment of LY34-bound mGluR3 structures revealed differences in inter-LBD and inter-CRD organization. Most notably, LY34 class 3 and LY34/VU602 class 2 show a large repositioning of LB2 and CRD compared to LY34 class 1. A smaller helix F/F angle (52.8°) is seen for LY34 class 1 compared to LY34 class 3 (58.3°) and LY34/VU602 (57.0°), indicative of the altered orientation between LB2 of adjacent subunits (**Fig. 6C**). Interestingly, the angle between helix B and helix O, which measures changes in inter-LB1 repositioning, is more similar for all three structures (97.7°, 101.4°, and 101.3° for LY34 class1, LY34 class3, and LY34/VU602, respectively), suggesting that most of the repositioning of the LBD between states takes place at the LB2 interface. CRD repositioning is also observed when measuring the structural offsets or the angle between CRDs when aligning the structures to LB1 of the same chain (**Fig. 6C; Fig. S8B**). Aligning all LY34-containing maps reveals that the CRD position for LY34 class 2 falls between that of class 1 and class 3, supporting the idea that class 2 represents a conformational intermediate (**Fig. S8C**). When aligning the maps obtained for LY34/VU602 class 1 and class 2, only a minor CRD offset is seen, suggesting that these two conformations are closely related and that PAM constrains the range of CRD motion (**Fig. S8D**). Unlike in our LY37-containing structures, all three LY34-containing models show very close single chain LBD alignment with an RMSD of 0.59 Å and 0.52 Å for LY34 class 3 vs LY34 class 1 or LY34/VU602 class 2, respectively. Additionally, we do not find any changes in LBD closure or LY34 binding site coordination regardless of the presence or absence of PAM (**Fig. S8E**). Together, these structures reveal that in the presence of LY34, the LBD samples conformations that are compatible with multiple CRD and TMD orientations, while the addition of PAM reduces the possible orientations sampled.

Alignment of intra-TMD conformation between LY34 bound class 1 and class 3 models shows an RMSD of 4.3 Å with modest repositioning of all helices (**Fig. S8F**). Alignment of LY34 class 3 with the LY34/VU602 structure reveals an improved alignment (RMSD= 3.0 Å), but with subtle outward shifts at the top of TM6 (**Fig. S8F**), which are consistent with the effects of PAM observed in agonist-bound structures. Consistent with this, the LY34/VU602 class 2 TMD also aligns more closely with the LY37/VU602 TMD (RMSD= 3.6 Å) compared to the LY37 only TMD (RMSD= 7.0 Å) (**Fig. S8F**) suggesting that VU602 stabilizes similar intra-TMD rearrangements independent of the orthosteric ligand.

The most dramatic differences between LY34-bound states are seen at the dimeric interface between the TMD bundles. The LY34 class 1 interface is asymmetric with the extracellular end of TM3 of one protomer and TM4 of the other protomer making contact (**Fig 6D, Fig. S8G**). The interface is not extensive due to bowing of TM4 and thus is tighter at the extracellular and intracellular ends. By aligning TMD bundles into the LY34 class 2 density (**Fig. S8H**), we can deduce that the inter-TMD interface of this state generally resembles that observed for class 1 but is slightly tighter and more symmetric such that the extracellular side of both TM3 helices contribute to the interface. The LY34 class 3 structure shows a strikingly different interface which only contains TM5 and extends throughout the length of the helix (**Fig. 6D; Fig. S8G**). A TM5 interface is also observed for LY34/VU602 class 2 (**Fig. 6D**) and is likely also present for LY34/VU602 class 1 (**Fig. S8I**). While the LY34/VU602 class 2 interface is similar to that observed with LY34 class 3, there is a slight reshaping throughout the length of the interface, such that they differ in interface distance at various points (**Fig. S8J**). Notably, this is the first tight TM5 containing interface observed for mGluR homodimers (**Fig. S8K, L**), although some prior mGluR homodimer structures show TM5 pointing towards each other at a distance that is too far to form interfacial contacts^30,38,44^. Furthermore, a recent cryo-EM study revealed a TM5 containing interface for an mGluR3-containing heterodimer^48^. Additionally, inactive GABA_B_ receptor structures also have been reported to contain variable TM5 interfaces ^70,74–76^ (**Fig. S8L**). Together these observations indicate that multiple family C GPCRs sample a TM5 containing interface but that the shape and stability of this interface is subtype and ligand dependent.

To further investigate the conformational rearrangements within our heterogeneous LY34 data sets, we performed 3DVA. When pooling the data either for LY34 class 1 and class 2 or LY34/VU602 class 1 and class 2, we observe LBD movements in both data sets leading to changes in LB2 angles (**Fig. S8M; supplemental Video 5,6)**. These angle changes are consistent with interconversion between the different helix F angles observed for our LY34 classes (**Fig. 6C**), which may accommodate the variety of CRD placements we observe across these structures (**Fig. S8B**) to accommodate both symmetric and asymmetric states with different TMD interfaces. We also observe a rotation and compaction movement for the CRDs as well as gain and loss of TMD density (**Fig. S8M**), indicating very dynamic TMDs. While the resolution is such that we cannot define specific changes to the TMD dimer interface, the loss and gain of density suggests that this is a highly dynamic area, sampling multiple conformations, and supporting the idea that we may be capturing a conformational intermediate along the activation pathway. We also performed 3DVA for individual classes of particles. For the LY34 class 3 data we observe TMD motion in which density assigned to TM4 moves towards the dimeric interface, suggesting the possibility of interconversions between a TM4 and TM5 containing interface (**Fig. S8M; supplemental Video 7)**. Together, these data support the finding that open, antagonist bound LBDs can accommodate multiple, inter-convertible CRD and TMD positions which can be restricted in the presence of a PAM.

### Toward a working model of mGluR activation, desensitization, and allosteric modulation

Together our structural analyses complement existing cryo-EM studies of mGluRs and allow a visualization of the complex conformational changes that occur across activation- and desensitization-associated states. Full-length, single-subunit alignments of the five structural models reported here reveal multiple classes of conformational shifts across domains (**Fig. 7A**). In brief, agonist binding stabilizes a closed LBD which leads to a large CRD repositioning such that LY34- and LY37-containing structures show very different TMD positions. Within each cluster of agonist and antagonist-bound structures, the presence or absence of PAM can fine-tune the position of the CRD and TMD. Repositioning of the CRD and TMD under different ligand-associated states leads to a range of inter-TMD dimer interfaces that exists along a corridor on the TMD surface starting with TM3 and TM4 (antagonist-bound), converting to a TM5 interface (antagonist-bound) and converging on a TM6 interface (agonist-bound) (**Fig. 7B**). PAMs appear to stabilize and subtly reshape the TM5 or TM6 interface in the presence of antagonist or agonist, respectively.

**Figure 7.**
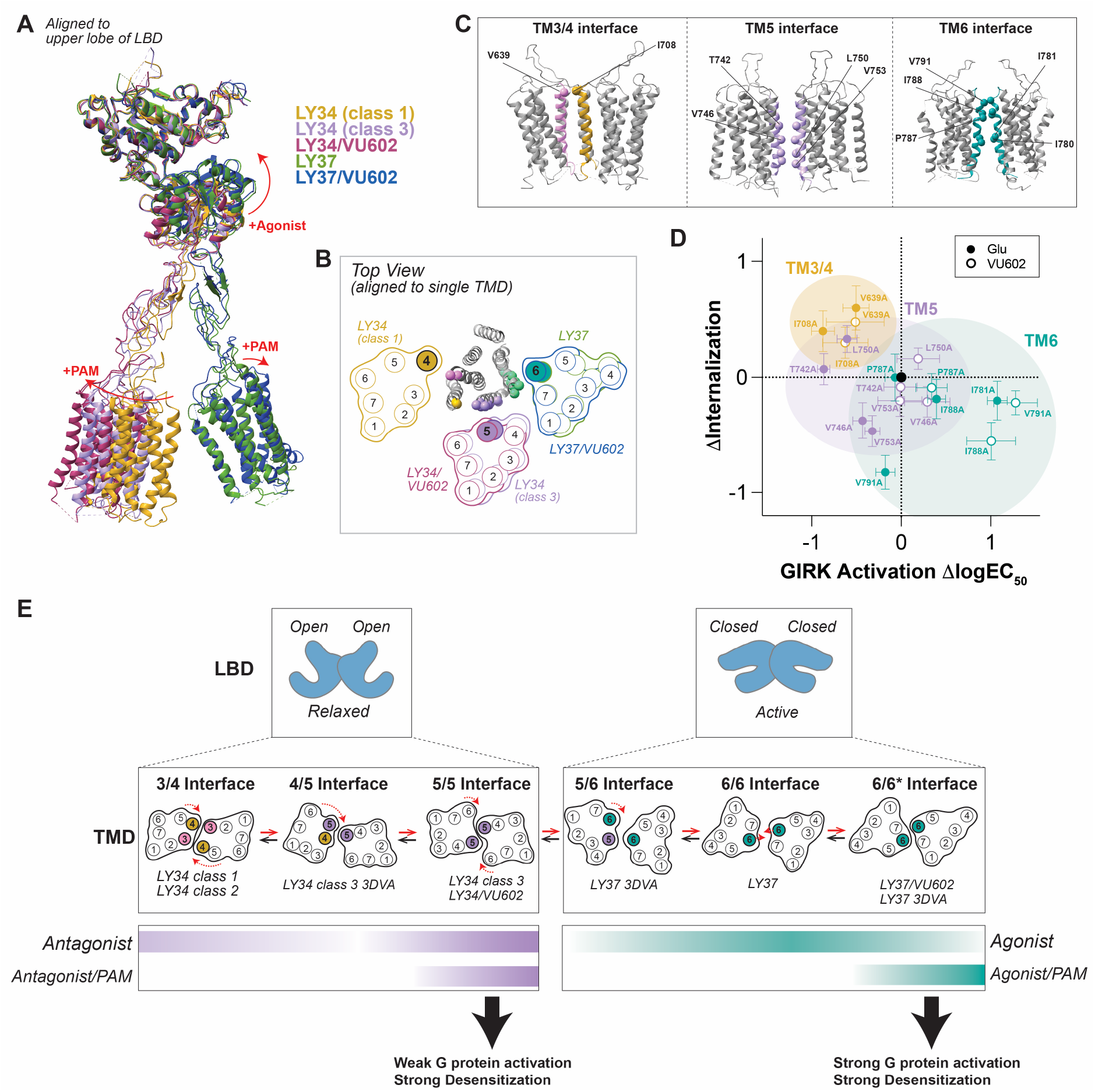
A rolling TMD interface working model of mGluR3 activation, allosteric modulation, and desensitization. **(A)** Single chain alignment for five mGluR3 structures aligned to LB1 of chain A with key rearrangements indicated via red arrows. **(B)** Top view of TMD interfaces when aligned to the TMD of chain A with LY34/VU602 TMD shown. Interfacial residues within 11 Å are shown as spheres. **(C)** TMDs of LY34 class1 (left), LY34/VU602 class 2 (middle), and LY37/VU602 (right) with the interface and key residues highlighted. **(D)** Scatter plot of the effects of alanine mutation to intersubunit interface interacting residues in terms of the response to glutamate or VU602 in G protein activation (x-axis) or internalization (y-axis) assays. Error bars represent the standard error of the mean of the difference between the mutant and the WT for Δlog EC50 or the maximum internalization difference. **(E)** Working model of a ligand-stabilized rolling TMD interface mechanism of activation. See also Figure S9.

To test the role of the identified TMD interfaces, we performed alanine scanning mutagenesis of interfacial residues observed in our structures (**Fig. 7C**). All alanine mutants expressed to normal levels (**Fig S9A, F, K**) and were tested in response to either glutamate or VU602 in both the G protein-dependent GIRK current assay and in the surface labeling internalization assay to assess both activation and desensitization (**Fig. 7D**). Mutation of residues V639 (TM3) and I708 (TM4) in the asymmetric TM3-TM4 interface had no effect on surface expression (**Fig. S9A**) but led to left-shifts in both agonist and PAM-evoked GIRK current dose-response curves (**Fig. S9B, C**) as well as an increase in the extent of internalization following treatment with either ligand (**Fig. S9D, E**). These results are consistent with the TM3-TM4 interface stabilizing inactive states that neither couple to G proteins nor promote GRK and *β*-arr-dependent endocytosis.

Mutations to interfacial TM5 residues had more complex results with comparable expression for all constructs (**Fig. S9F**) but divergent responses to ligands (**Fig. 7D**). Glutamate-driven GIRK activation was left-shifted for all mutants (**Fig. S9G**) while PAM-driven GIRK currents were unaffected, except for L750A which showed a reduced maximal response (**Fig. S9H**). In response to glutamate, all variants showed clear internalization with a substantial and modest enhancement, respectively, for L750A and T742A. In contrast, V746A and V753A showed impaired internalization (**Fig. S9I**). The same trends were observed in PAM induced internalization but with more subtle effects (**Fig. S9J**). It’s notable that in response to glutamate V746A and V753A showed enhanced G protein activation and impaired GRK/*β*-arr coupling (**Fig. 7C**), suggesting the presence of biased conformations with preferential G protein-coupling. Together, these data show diversity in the role of TM5 residues in shaping the ligand response, revealing underexplored complexity in the conformational states that mediate mGluR activation and desensitization.

Prior work has shown that the highly conserved TM6 interface is involved in activation of the mGluR receptors, as showed by cross-linking or by mutating residues at the extracellular end of the helix^38,56,77^. We extended this analysis by mutating TM6 residues along the extended interface that is re-shaped in the presence of PAM (**Fig. 5F-H; S6I**). All mutants expressed to comparable levels (**Fig. S9K**) and showed glutamate- and PAM-evoked GIRK current dose-response curves that were either wild-type like or right-shifted (**Fig. S9L, M**). In the most extreme case, I780A showed no response to either ligand (**Fig. S9L, M**) and I781A showed no response to PAM. Consistent with impaired G protein activation seen with the GIRK current assay, all TM6 mutants either had no effect or decreased internalization in response to glutamate or PAM (**Fig. S9N, O**). Notably, the most extracellular residue mutated on TM6, V791A, showed a much larger impairment of internalization in response to glutamate than PAM (**Fig. 7D**). As was seen with GIRK currents, I780A showed no internalization in response to glutamate or PAM (**Fig. S9N, O**) while I781A and I788A impaired only PAM-driven internalization. These data point to the outward facing surface of TM6, particularly residues at the mid-point of the helix, as a key site for regulating mGluR activation and desensitization.

While our functional data support the importance of TM5 and TM6 interfaces, it is important to note that TM5 and TM6 are both known to contain key residues that form PAM binding sites^28,45,46,78^ and critical molecular switches involved in intra-subunit TMD activation^30,50,79^. Thus, the functional effects of our mutants are likely due to a combination of effects on dimer interface strength/shape, intra-TMD conformational dynamics, and PAM binding. Despite this ambiguity, these data clearly highlight the inter-TMD interface as a hotspot for controlling the response to both PAMs and agonists in terms of both G protein-dependent activation and GRK/*β*-arr-dependent internalization.

Together these functional and structural data are in line with a working model of mGluR3 activation and desensitization based on a rolling, stepwise inter-TMD reorientation which is outlined in **Fig. 7E**. In the presence of antagonist-stabilized open-open/relaxed LBDs, we see an ensemble of TM3-, TM4-, and TM5-containing interfaces. With the addition of PAM, the TM5-containing interfaces become the dominant class. In the presence of agonist-stabilized closed-closed/active LBDs, a variety of TM6-containing interfaces are occupied, including an asymmetric TM5/TM6 interface observed with 3DVA, with PAM ultimately stabilizing the re-shaped “6/6*” interface with tighter interactions at the extracellular end of TM6.

## Discussion

Our results reveal numerous functional features of mGluR allosteric modulation and provide a structural framework for their interpretation. We find that mGluRs can both activate and desensitize in response to orthosteric agonists and PAMs, although PAMs typically show a bias toward desensitization. Furthermore, application of PAMs can increase the maximum response to agonists, although this is receptor subtype and ligand specific. The complex ligand responses of mGluRs are underscored by conformational heterogeneity, where a variety of LBD and TMD conformations can be differentially stabilized in various states, as observed in our full-length cryo-EM structures of mGluR3 and prior structural analyses^30,38,44–48^. Together these results have a plethora of implications regarding both the biophysical mechanisms and physiological properties of mGluRs.

### Biophysical/Structural implications

The ability of PAMs to both serve as modulators of orthosteric agonists as well as direct allosteric agonists, even in the presence of orthosteric antagonists, is consistent with a variety of inactive and active conformations that are differentially populated depending on ligand occupancy. Indeed, our structural data supports a model where the activation pathway passes through multiple TMD interfaces including the hydrophobic outer surface of TM3, TM4, TM5 and TM6 (**Fig. 7**). This model posits that these inter-TMD interfaces are highly sensitive to ligand conditions and rapidly interchangeable, which may be facilitated by the hydrophobic and limited interaction surface of the dimer interfaces observed by cryo-EM. Our model is supported by a seminal biochemical crosslinking study of mGluR2 which showed ligand-sensitive crosslinking of outward facing residues in TM4, TM5, and TM6^77^.

In line with both this prior crosslinking study and published cryo-EM structures, the agonist LY37 stabilizes a closed-closed/active LBD dimer conformation and an inter-TM6 interface in mGluR3 (**Fig. 5**). Interestingly, the addition of the PAM VU602 stabilizes a very similar state but with some key differences which likely explain the ability of PAMs to enhance agonist affinity and efficacy. At the LBD and CRD level, VU602 leads to a tightening of LBD and CRD interfaces along with a compaction of the orthosteric binding site such that closer interactions are made with the agonist. At the TMDs, PAMs stabilize a re-positioned TM6 interface including interactions at the extracellular end of the helix. Critically, this state appears to be populated in the absence of VU602 in 3DVA of the LY37 only data but is likely only fully stabilized by PAM and/or G protein transducer^55^. This is consistent with a conformational selection model of allosteric modulation and our observation that PAMs increase the maximal inter-TMD FRET response to agonist (**Fig. 4**). Why does the TM6 interface play such a critical role in mGluR activation? One possible explanation is the ability of this interface to influence the conformation of the previously identified tryptophan “toggle switch” (W782 in mGluR3). This conserved tryptophan is thought to act as a rotameric microswitch that controls the movement and compaction of TM3, TM5, and TM6 to promote the inter-helical reorientations required for activation across GPCR families^30,50,51,78–80^. Small changes in TM6 positioning due to TMD interface shape may thus reposition the helix in a way that allows for the toggle switch to stabilize an active state. In line with this, subtle mutation (I780A, I781A) to interfacial residues adjacent to the toggle switch severely impaired or abolished mGluR3 activation and internalization.

Our data also revealed that in the presence of open LBDs stabilized by an orthosteric antagonist, PAMs can still exert functionally relevant effects. We show that a PAM can drive strong mGluR3 internalization, but not G protein activation, in the presence of an orthosteric antagonist (**Fig. 4**). In contrast, we find that mGluR2 is capable of PAM-driven G protein activation even in the presence of an orthosteric antagonist, consistent with our previous studies^29^ and what has also been seen in mGluR4^28^ and mGluR5^30^. This reveals mGluR subtype differences in PAM-evoked responses, which may be driven by subtype-specific conformational and structural properties as previously observed in smFRET and TMD dimerization studies^41,56,81^. Our cryo-EM structures of mGluR3 in the presence of the antagonist LY34 reveal dramatic structural variability with three primary states observed containing TM3/4 (class 1, class 2) or TM5 (class 3) inter-TMD interfaces and variable inter-LB2 and CRD orientations. Consistent with the ability of PAM to drive inter-TMD FRET changes even in the presence of LY34 (**Fig. 4**), the addition of PAM stabilized a TM5 interface-containing states with similarity to the class 3 state observed with LY34 alone. This TM5 interface represents a unique inter-TMD interface among homodimeric mGluR structures, although a recent mGluR2/3 heterodimer cryo-EM study observed similar TM5 interfaces under various antagonist-stabilized conditions^48^. Notably, an antagonist-insensitive component of the PAM conformational response was also seen in mGluR2 in a recent inter-CRD and inter-TMD FRET study^35^, suggesting that PAM-stabilized intermediates are a common feature of mGluRs. As was also seen in the agonist-bound structures, in our LY34-bound structures VU602 also modified the TMD conformation by re-shaping the TM5 interface and driving an opening of the extracellular face of the TMD.

Together our work raises the question of why PAMs show an apparent desensitization bias compared to agonists. Based on our structural and functional data we posit that the distinct TMD interface stabilized by a given ligand or ligand combination controls transducer coupling in two ways. First, different TMD interfaces may provide steric constrains that determine the ability of transducers (G proteins, GRKs, *β*-arrs) to access their binding sites on the intracellular face of the receptor. Indeed, in prior analysis of a TM4-containing interface of mGluR2, Seven et al proposed that this conformation was incompatible with G protein coupling to either subunit due to steric clashes with either the other subunit or the plasma membrane^46^. Second, different TMD interfaces may stabilize distinct intra-TMD conformations with intrinsic preferences for G proteins, GRKs, or *β*-arrs, as has been proposed for family A GPCRs^12–15^ Based on our functional data where the antagonist/PAM ligand combination drives strong internalization, but not G protein-coupling, we reason that the TM5 interface states we observe are intrinsically *β*-arr biased (**Fig. 7E**) either due to steric barriers to G protein heterotrimer coupling, incompatible intra-TMD conformations, or both. We speculate that GRKs and *β*-arrs may be less constrained by the TMD interface given that their key interactions are with flexible C-terminal domains. Indeed, *β*-arrs have been proposed to couple to GPCRs in “tail-only” modes where TMD core interactions are not required^13,82,83^. Furthermore, *β*-arrs have been shown to bind GPCR TMD cores with variable angles^84,85^. It is important to note that in conditions where PAM is applied in the absence of agonist or antagonist it is likely that a more complex combination of conformations, including intermediate states not observed in cryo-EM, are occupied. This likely includes TM6 interfaces as PAM-driven GIRK activation and internalization are both sensitive to TM6 interface mutations (**Fig. 7D**). Presumably, the ability of PAM to stabilize both TM5 (“desensitization-biased”) and TM6 (“balanced”) interfaces enables a net bias toward GRK and *β*-arr mediated desensitization processes. In the presence of agonists, PAMs can enhance both G protein activation and desensitization, consistent with the TM6 interface composing a state or ensemble of states with strong coupling to both G protein and GRK/*β*-arrs (**Fig. 7E**), although the precise conformations involved are not clear. Ultimately, further work, including higher resolution structural data, is needed to understand how the inter-TMD interface controls intra-TMD rearrangements and G protein versus *β*-arr coupling.

### Physiological/Therapeutic implications

The myriad effects of PAMs on both mGluR activation and desensitization complicates interpretation of pharmacological studies and therapeutic application of such ligands. In the case of mGluR3, preclinical studies have pointed to specific activation of this subtype as a promising strategy for the treatment of cognitive deficits associated with neuropsychiatric diseases, including schizophrenia^86–88^, for neuroprotective effects in the context of neurodegenerative diseases^89–91^, and for the treatment of addiction^92^. Such studies have motivated the ongoing development of subtype-specific mGluR3 PAMs^63^. To date, VU602 is the only reported compound with PAM activity toward mGluR3 although it is also active toward mGluR2.

Our recent finding of strong mGluR3 coupling with GRKs and *β*-arrs^57^, along with new findings reported here showing enhanced desensitization with VU602, suggests previously unappreciated complexity in the pharmacological targeting of this receptor and other mGluRs. Depending on the context, the strong GRK/*β*-arr coupling driven by PAMs may limit or enhance their therapeutic effects relative to orthosteric agonists. For example, PAMs may boost signaling in the short term but drive a long-term down regulation of mGluRs which limits and leads to tolerance. Alternatively, the enhanced *β*-arr coupling of PAMs may initiate signaling pathways that drive the beneficial effects of mGluR activation, as has been proposed for some family A GPCR agonists^59,60^. Our findings that PAMs are also desensitization-biased for mGluR7 and mGluR8 and can even drive the internalization of mGluR2, an otherwise desensitization-resistant subtype, suggests that this is a critical aspect of allosteric pharmacology to be considered for all mGluR-targeting compounds. Furthermore, the complexity of TMD conformational changes revealed by our work, suggests that it may be possible to develop *β*-arr or G protein-biased allosteric compounds for mGluRs. Overall, extensive future work is needed to better understand the effects of G protein-dependent and independent signaling and trafficking of mGluRs in neurological and behavioral contexts and to harness this for improved therapeutic approaches.

### Limitations of the Study

As in most structural studies of GPCRs, our cryo-EM analysis is based on snapshots of detergent-solubilized receptors, raising the questions of how well this *in vitro* system correlates to the structural states and dynamics of mGluRs in their physiological membrane environments. Furthermore, the resolution of our structures, particularly in the TMDs, limits our ability to draw strong conclusions about some aspects of ligand-associated structural changes. For example, we did not resolve sufficient density to identify PAM binding sites, including resolving if one or two PAMs bind within an mGluR3 dimer. Furthermore, we chose to constrain the structural dynamics of mGluR3 by performing cryo-EM in the presence of either saturating agonist or antagonist, rather than in the absence of either. Structural analysis with PAM alone may reveal novel states along the activation pathway and provide further insight into the unique activation and desensitization properties of PAMs. Along these lines, our cryo-EM analysis is likely to miss rarely populated but potentially key intermediate states. This could include states where TM6 interfaces are occupied in the presence of antagonist, although no evidence for such a state is seen in 3DVA. Finally, all functional experiments reported in this study were performed in HEK 293 cells. While these cells provide an excellent platform for assessing basic receptor properties, it will be critical in future work to extend these studies to the biological context of neurons and glial cells, especially in the highly organized synaptic compartments where mGluRs typically function^16,93^. Overall, despite these limitations this study reveals new and key dimensions of mGluR structure, function, and pharmacology that should drive future work on these complex receptors and other family C GPCRs.

## Methods

### Molecular biology

Rat SNAP-mGluR2, SNAP-mGluR3, SNAP-mGluR7 and human SNAP-mGluR8 constructs used in this study were previously generated^22,57^. The SNAP-mGluR3-Halo-ICL2 FRET sensors was previously reported^56^. All SNAP-mGluR constructs contain an N-terminal signal sequence from rat mGluR5 followed by an HA epitope and the SNAP-tag self-labelling enzyme upstream of the mGluR sequence and are inserted in a pRK5 vector backbone. Single point mutagenesis was performed using PCR-mediated site-directed mutagenesis. Previously described GIRK1-F137S homotetramerization mutant^94^ and tdTomato (Addgene #30530) were used for patch-clamp experiments. GRK2-GFP plasmid^5^ was used in internalization and acute desensitization experiments with mGluR2. Dominant negative β-arr1 (S412D) construct^57,95^ was used to confirm clathrin dependent internalization of mGluR3 in response to agonist or PAM. Glutamate transporter (EAAT1) plasmid was kindly gifted by Prof. Miriam Stoeber (University of Geneva, Switzerland). For cryo-EM studies, full length rat mGluR3 with a C-terminal thrombin cleavage site followed by a C-terminal mVenus and Twin-Strep-tag^96^ was generated in the pEZT-BM BacMam expression vector^97^.

### Cell culture and transfection

HEK293 (ATCC: CRL-1573) and HEK293T (ATCC: CRL-3216) cells were grown and maintained in Dulbecco’s Modified Eagle’s Medium (DMEM; #10-013-CV Corning, USA) supplemented with 10% fetal bovine serum (FBS; Thermo Fisher #A5256) and cultured at 37°C/5% CO_2_. HEK293 GRK2/3 knock-out cells were kindly gifted by Asuka Inoue (Tohoku University, Japan). For electrophysiology and imaging experiments, cells were seeded on poly-L-lysine (Sigma #P2636) coated rounded 18 mm glass coverslips and transfected using Lipofectamine 2000 (Thermo Fisher Scientific). 6 hr after transfection media was replaced and supplemented with antagonist until the time of the experiment (5 µM LY34 for mGluR2 and mGluR3, 20 µM CPPG for mGluR7 and mGluR8) to maintain cell health. Experiments were performed 24-48 hr post-transfection.

### Compound Synthesis and Medicinal Chemistry

Detailed in vitro evaluation of VU602 and structurally related compounds can be found in Yamada et al^63^.

#### General

All NMR spectra were recorded on a 400 MHz AMX Bruker NMR spectrometer. 1H and 13C chemical shifts are reported in δ values in ppm downfield with the deuterated solvent as the internal standard. Data are reported as follows: chemical shift, multiplicity (s = singlet, d = doublet, t = triplet, q = quartet, b = broad, m = multiplet), integration, coupling constant (Hz). Low resolution mass spectra were obtained on an Agilent 6120 or 6150 with ESI source. Method A: MS parameters were as follows: fragmentor: 70, capillary voltage: 3000 V, nebulizer pressure: 30 psig, drying gas flow: 13 L/min, drying gas temperature: 350 °C. Samples were introduced via an Agilent 1290 UHPLC comprised of a G4220A binary pump, G4226A ALS, G1316C TCC, and G4212A DAD with ULD flow cell. UV absorption was generally observed at 215 nm and 254 nm with a 4 nm bandwidth. Column: Waters Acquity BEH C18, 1.0 x 50 mm, 1.7 um. Gradient conditions: 5% to 95% CH3CN in H2O (0.1% TFA) over 1.4 min, hold at 95% CH3CN for 0.1 min, 0.5 mL/min, 55 °C. Method B: MS parameters were as follows: fragmentor: 100, capillary voltage: 3000 V, nebulizer pressure: 40 psig, drying gas flow: 11 L/min, drying gas temperature: 350 °C. Samples were introduced via an Agilent 1200 HPLC comprised of a degasser, G1312A binary pump, G1367B HP-ALS, G1316A TCC, G1315D DAD, and a Varian 380 ELSD (if applicable). UV absorption was generally observed at 215 nm and 254 nm with a 4 nm bandwidth. Column: Thermo Accucore C18, 2.1 x 30 mm, 2.6 um. Gradient conditions: 7% to 95% CH3CN in H2O (0.1% TFA) over 1.6 min, hold at 95% CH3CN for 0.35 min, 1.5 mL/min, 45 °C. High resolution mass spectra were obtained on an Agilent 6540 UHD Q-TOF with ESI source. MS parameters were as follows: fragmentor: 150, capillary voltage: 3500 V, nebulizer pressure: 60 psig, drying gas flow: 13 L/min, drying gas temperature: 275 °C. Samples were introduced via an Agilent 1200 UHPLC comprised of a G4220A binary pump, G4226A 3 ALS, G1316C TCC, and G4212A DAD with ULD flow cell. UV absorption was observed at 215 nm and 254 nm with a 4 nm bandwidth. Column: Agilent Zorbax Extend C18, 1.8 µm, 2.1 x 50 mm. Gradient conditions: 5% to 95% CH3CN in H2O (0.1% formic acid) over 1 min, hold at 95% CH3CN for 0.1 min, 0.5 mL/min, 40 °C. Optical specific rotations were obtained using JASCO P-2000 Digital Polarimeter equipped with Tungsten-Halogen lamp (WI), 589 nm wavelength, photomultiplier tube (1P28-01) detector and CG2-100 Cylindrical glass cell, 2.5ø x 100 mm. For compounds that were purified on a Gilson preparative reversed-phase HPLC, the system comprised of a 333 aqueous pump with solvent-selection valve, 334 organic pump, GX271 or GX-281 liquid hander, two column switching valves, and a 155 UV detector. UV wavelength for fraction collection was user-defined, with absorbance at 254 nm always monitored. Method 1: Phenomenex Axia-packed Luna C18, 30 x 50 mm, 5 µm column. Mobile phase: CH3CN in H2O (0.1% TFA). Gradient conditions: 0.75 min equilibration, followed by user defined gradient (starting organic percentage, ending organic percentage, duration), hold at 95% CH3CN in H2O (0.1% TFA) for 1 min, 50 mL/min, 23 °C. Method 2: Phenomenex Axia packed Gemini C18, 50 x 250 mm, 10 um column. Mobile phase: CH3CN in H2O (0.1% TFA). Gradient conditions: 7 min equilibration, followed by user defined gradient (starting organic percentage, ending organic percentage, duration), hold at 95% CH3CN in H2O (0.1% TFA) for 7 min, 120 mL/min, 23 °C. Chiral separation was performed on a Thar (Waters) Investigator SFC Column: Chiral Technologies CHIRALPAK IF, 4.6 x 250 mm, 5 µm column. Gradient conditions: 20% to 50% IPA in CO2 over 7 min, hold at 50% CO2 for 1 min. Flow rate: 3.5 mL/min. Column temperature: 40 °C. System backpressure: 100 bar. Solvents for extraction, washing and chromatography were HPLC grade. All reagents were purchased from Aldrich Chemical Co. and were used without purification.

#### Preparation of (R)-3-(3-(4-(3,3-dimethylbutanoyl)-3-hydroxy-2-methylphenoxy)-2-methylpropoxy)-4-methoxybenzoic acid (VU602)

In a microwave vial equipped with a stir bar, (*S*)-1-(2-hydroxy-4-(3-hydroxy-2-methylpropoxy)-3-methylphenyl)-3,3-dimethylbutan-1-one (41 mg, 0.14 mmol), methyl vanillate (13 mg, 0.07 mmol), di-tert-butyl azodicarboxylate (35 mg, 0.15 mmol) and triphenylphosphine (40 mg, 0.15 mmol) in THF (0.5 mL). The reaction was heated to 120 °C for 30 minutes in a Biotage microwave reactor. After completion, the reaction was diluted with water and the desired product extracted with dichloromethane (3X) and the organic layer passed through a phase separator. The product was concentrated and purified using reverse phase HPLC: 30x100mm column, 35-100% acetonitrile gradient (w/ 0.1% TFA) over 10 minutes. Fractions containing the desired product were concentrated to give compound the title compound: **VU602** (29% yield), HR-MS observed m/z = 445.2228; calculated m/z = 445.2221. LC-MS Retention time: 1.638 min, [M+H]^+^: 445.2. 1H NMR (400 MHz, CDCl3) δ 7.68 (dd, J = 8.4, 2.0 Hz, 1H), 7.57 – 7.51 (m, 2H), 6.88 (d, J = 8.5 Hz, 1H), 6.39 (d, J = 9.0 Hz, 1H), 4.17 – 4.00 (m, 4H), 3.84 (s, 3H), 2.71 (s, 2H), 2.54 (s, 1H), 2.04 (s, 3H), 1.18 (d, J = 6.9 Hz, 3H), 1.00 (s, 9H).

### Patch-clamp electrophysiology

Whole-cell patch-clamp recordings of cells co-expressing mGluRs with GIRK1-F137S and tdTomato were performed 24-36 hr post-transfection as previously described^29,56^. GIRK currents were recorded at -60 mV in voltage clamp mode using an Axopatch 200B amplifier and a Digidata 1550B interface commanded via pClampex software (Molecular Devices). Bath solution contained (in mM): 120 KCl, 25 NaCl, 10 HEPES, 2 CaCl_2_, 1 MgCl_2_. In this bath solution, different agonists, antagonists, or allosteric modulators were added via gravity-driven perfusion at the concentrations specified for each experiment. For all experiments, drug responses were normalized to glutamate as all recordings concluded with application of saturating glutamate (100 μM for mGluR3; 1 mM for mGluR2, mGluR8; 10 mM for mGluR7). All data was obtained from at least two separate experimental days for each condition. Each condition was measured from at least 3 different cells. For comparison of EC_50_ shifts of the different mutants in **Fig. S9**, statistical analysis was performed using an F-test where the null hypothesis was the equal distribution of all mutant dose response curves in comparison to the WT.

### Widefield fluorescence imaging and surface labelling internalization assay

To visualize internalization, HEK293T cells were transfected with 0.7 µg of SNAP-tagged receptors per well of a 12-well plate. To reduce ambient glutamate levels and prevent basal internalization in the mGluR3 conditions, cells were co-transfected with 0.7 µg of the glutamate transporter (EAAT1). 24 hr post-transfection, cells were labeled for 30 min at 37° C with 1 µM Surface BG-Alexa546 (impermeable dye; New England Biolabs) for SNAP in extracellular buffer (EX) containing (in mM): 10 HEPES, 135 NaCl, 5.4 KCl, 2 CaCl_2_, 1 MgCl_2_ and pH 7.4. After labelling, cells were thoroughly washed with fresh EX buffer and incubated in agonist/antagonist and/or PAM/NAM for 30 min. Live cells were imaged on an Olympus IX83 inverted microscope through an CMOS ORCAScience Flash4v3.0 camera (Hamamatsu) and using a 100X immersion oil objective (NA 1.45). Fluorophore was excited using a coupled 561 nm laser. A minimum of three days of independent transfections and a minimum of 7 images were acquired per condition. For the surface labelling assay, we followed a previously standardized protocol^6,57^. Briefly, after 24 hr expression, SNAP-tagged receptors were incubated in prewarmed culture media containing the desired combination of agonist, antagonist, PAM and/or NAM for 60 min at 37° C. Afterwards, cells were labelled with 1 µM Surface BG-Alexa 546 in EX for 20 min at RT and imaged on an Olympus IX83 microscope using a 60X immersion oil objective (NA 1.49). 10-12 snapshots per condition were acquired per day and a minimum of three independent replicates on different days were used per condition. Quantification of total membrane fluorescence of cells was done using a Li algorithm threshold method in Image J (Fiji). The mean intensity of cells was calculated by subtracting the background for each condition and normalizing against the control antagonist condition. mGluR2 internalization experiments (**Fig. 3K, L**) were performed with co-expression of wild type GRK2.

### Ensemble FRET measurements

Live cell ensemble, intersubunit FRET measurements were performed as previously described^56^. Briefly, 36-48 hr post-transfection, cells were washed with EX buffer, labeled with fluorophores for 45 min at 37 ° C, and washed again in EX. For inter-LBD FRET experiments, cells expressing N-terminal SNAP-tagged mGluRs were labeled with 1 µM BG-LD555 (Lumidyne Technologies) and 3 µM BG-LD655 (Lumidyne Technologies). For inter-TMD FRET experiments, cells expressing mGluR3 with Halo tag in ICL2 were labeled with 1 µM CA-JF549 (Promega) and 3 µM CA-JF646 (Promega). Cells were then imaged at room temperature with a 60x objective (N.A. = 1.45) on an inverted Olympus IX83 microscope with donor excitation via a 561 nm laser and imaging of both donor and acceptor channels at 0.5-1 Hz with an exposure time of 100 ms using two separate sCMOS cameras (Hamamatsu ORCA-Flash4v3.0). A gravity-driven perfusion system was used to deliver drugs diluted in EX solution. Fluorescence intensity traces were extracted from each movie by drawing a region of interest (ROI) around a cell cluster (3-15 cells) from donor and acceptor channels using CellSens software (Olympus). FRET was calculated as FRET = I_Acceptor_ / (I_Donor_ + I_Acceptor_), in which I = fluorescence intensity and the FRET value was normalized based on the initial baseline before any drug application. For all experiments, data was obtained from multiple cell clusters and averaged from at least three replicates from at least two separate transfections.

### Expression and purification of mGluR3

cDNA for rat mGluR3 was cloned into the pEZT-BM BacMam expression vector^97^ and the C-terminus was fused to a thrombin cleavage site preceding a C-terminal mVenus and Twin-Strep-tag^96^. The expression construct was transformed into DH10Bac cells to produce bacmid which was then transfected into *Spodoptera frugiperda* (Sf9) cells grown in ESF 921 media (Expression Systems). P1 and P2 virus production was monitored using GFP fluorescence from the pEZT-BM vector until virus harvesting. HEK293S GnTI^-^ cells (ATCC CRL-3022) were grown at 37 °C and 8% CO_2_ to a density of 3.5 × 10^6^ cells/mL in FreeStyle suspension media (Gibco) supplemented with 2% fetal bovine serum (Gibco) and Anti-Anti (Gibco). P2 virus was added to cells, the suspension was incubated at 37 °C for 24 h, then sodium butyrate (Sigma) was added to a final concentration of 10 mM and flasks were shifted to 30 °C and 8% CO2. Cells were collected 72 hours after transduction by low-speed centrifugation, flash-frozen in liquid nitrogen, and stored at −80 °C.

Cell pellets were resuspended in cold lysis buffer containing 10 mM HEPES (pH 7.5),10 mM MgCl2, 20 mM KCl, 1 M mM NaCl, EDTA-free protease inhibitor cocktail tablet (Sigma), and 25 ug/ml DNase (50 mL buffer per ∼1g of cell pellet). Following manual pipetting, the cells were disrupted using a QSonica Q700 sonicator (5 × 15 sec, power level 60). The lysate was then clarified by low-speed centrifugation at 7,200 × *g* for 20 minutes and membranes were isolated by ultracentrifugation at 125,000 × *g* for 2 hours. All steps were performed on ice or at 4°C. Membranes were then collected and covered with 1mL membrane freezing buffer containing 10 mM HEPES (pH 7.5),10 mM MgCl2, 20 mM KCl, and 30% glycerol, flash-frozen in liquid nitrogen, and stored at −80 °C. For both LY34 bound structures, 20 *μ*M LY34 was present in all steps.

Membranes were resuspended and Dounce homogenized in buffer containing 100 mM HEPES (pH 7.5), 300 mM NaCl, 2 mM CaCl2, and the appropriate ligand (10 *μ*M LY37, 20 *μ*M LY34, 10 *μ*M VU602). For the LY34 plus VU602 preparation only, the sample was incubated with an additional 100 *μ*M VU602 for 30 minutes on ice prior to proceeding to the next step. Protein was extracted via the addition of n-Dodecyl-β-D-maltopyranoside (DDM) with cholesteryl hemisuccinate (CHS) (Anatrace, D310-CH210) to a final concentration of 50 mM DDM / 4 mM CHS and rotated for 1 hour at 4 °C. Following ultracentrifugation at 4°C at 125,000 × *g* for 50 min the supernatant was filtered and bound to a 5 mL Strep-Tactin column (GE) equilibrated with running buffer containing 100 mM HEPES (pH 7.5), 300 mM NaCl, 2 mM CaCl_2_, 0.1% glyco-diosgenin (GDN), and the appropriate ligand. To facilitate detergent exchange, the column was washed with running buffer for 5x column volumes. To remove bound heat shock proteins, the column was washed with 5x column volume of running buffer supplemented with 10 mM MgCl_2_ and 5 mM ATP and then 5x more washes of running buffer. Protein was subsequently eluted via running buffer supplemented with 5 mM D-Desthiobiotin (IBA). Elution fractions were collected, analyzed by SDS-PAGE and concentrated using 100KD cut-off. The sample was loaded onto a Superose 6 Increase 10/300 GL (Cytiva) column equilibrated in buffer containing 100 mM HEPES (pH 7.5), 300 mM NaCl, 2 mM CaCl2, 0.01% GDN, and the appropriate ligand. Elution fractions were collected, analyzed by SDS-PAGE, and key fractions were concentrated using a 100KDa cut off and used for cryo-EM studies.

### Cryo-EM sample preparation, data acquisition, and processing

mGluR3 samples ranging from 2.0-4.7 mg/mL were applied at a volume of 3 ***μ***L to plasma treated UltrAuFoil 1.2/1.3 300 mesh grids (Quantifoil). Vitrified samples were prepared using a Vitrobot Mk IV (Thermo Fisher), blot time of 2 seconds, blot force of -2, at 100% humidity and 23°C. Grids were imaged with a Titan Krios electron microscope (Thermo Fisher) operated at 300 kV and a nominal magnification of 105,000× and equipped with a K3 camera (Gatan) set in super-resolution mode (0.426 Å pixel size). Images were collected using Leginon.

Beam induced motion was corrected using MotionCor2 in Relion 3.1^98^ using two-fold binning and dose-weighting. These images were then used for contrast transfer function estimation via CTFFIND4.1^99^. In Relion, particles were picked using a 3D initial model template of the receptor and extracted using a box size of 416. Further processing was then performed in CryoSPARC version 3.3.2^100^. Ab initio followed by multiple rounds of heterogeneous refinement resulted in one good class that was further refined via subsequent ab initio and heterogeneous refinement jobs (**Fig. S7, S9**). The best heterogeneous refinement class/classes were then used for homogeneous refinement followed by non-uniform refinement (C2 for LY37, LY37/VU602, LY34/VU602, and C1 for LY34). These particles were then imported back into Relion for Bayesian polishing^101^ followed by non-uniform refinement to obtain the best full length map. Using a mask of the extracellular domain (LBDs and CRDs), local refinement was performed to obtain a map of the extracellular domain at improved resolution. Additionally using a mask of the TMD, particle subtraction was performed to generate a particle stack only containing TMD information followed by 2D classification to exclude a small number of poorly resolved particles, and then an ab initio job to generate a 3D volume and non-uniform refinement. This results in a map of the TMD with improved density for helices that are not well resolved in the full-length map. When applicable, 3D variability analysis was performed on full-length maps with a mask that excluded solvent and micelle, with 5 variability components and a low-pass filter resolution of 5 Å.

### Model building

To ensure proper alignment of maps, the ECD and TMD maps were aligned to their respective full-length map. Initial models were built using mGluR3 homology models for 7MTR (used for LY37 and LY37/VU602), 7EPA (used for LY34 class 1), 7WI6 (used for LY34/VU602) or our own previous models. Each model was then refined via iterative rounds of real-space refinement using the full length map in Phenix^102^ and manual refinement in Coot^103^ using information from both the full length and domain maps. Regions with poor resolution were either deleted, as in the case of some loops, or stubbed, as is the case of the TMDs. Visual representations were prepared using ChimeraX^104^ and Pymol^105^.

To prepare the models for cavity analysis, the missing atomic coordinates in chain A of the LY37 and LY37/VU602 cryo-EM structures without N- and C-termini (residues 31-822) were generated using the MODELLER software version 9.19^106^ and the rat mGluR3 sequence (UniProt ID P31422). Specifically, missing segments from the extracellular domain (ECD; residues 120-139) and intracellular loop 2 (IC2; residues 664-685) were modeled based on the mGluR2 structures corresponding to PDB IDs 7EPB and 7E9G, respectively, while missing TMD side chains and segments from intracellular loop 1 (IC1; residues 600-610), intracellular loop 3 (IC3; residues 759-766), and extracellular loop 3 (EC3; residues 794-800) were based on the mGluR2 structure corresponding to PDB ID 7MTR. All remaining segments with fewer than four residues were built *ab initio*. Reconstructed models of chain A for each structure were selected based on the lowest *Discrete Optimized Protein Energy* (DOPE) score^107^, and then copied and overlapped onto chain B using the PyMOL Molecular Graphics System (version 2.5.2). The final models were prepared for analysis using the Protein Preparation Wizard within the Schrödinger Maestro software package^108^ and the default protocol. The latter involved adding missing side chains and hydrogen atoms, capping the two protomers at residues R31 and K822, assigning correct bond orders and side chain protonation states at pH 7.4, and performing a short, restrained minimization *in vacuo* using the OPLS3e force field^109^.

### Cavity analysis

MOL2 files for chain A of the LY37 and LY37/PAM structures were generated using ChimeraX^110^. The IChem toolkit (version 5.2.9)^111,112^ was used to perform a cavity analysis on each structure. Specifically, the cavity detection and druggability/ligandability prediction tool called Volsite was used in unrestricted mode. The identified cavities were then assessed based on a measured druggability score (drugg score) with positive values indicating druggable cavities and negative values indicating non-druggable cavities. Molecular graphics were created using PyMOL.

### Quantification and Statistical Analysis

Statistical analysis for all experiments is indicated in figure legends, with error bars representing standard error of the mean (SEM) and each data point being repeated in triplicate or more. Analysis was performed using GraphPad Prism 9.0.

### Data availability

The atomic coordinates and cryo-EM density maps for mGluR3 bound to LY37, LY37/VU602, LY34 (class 1), LY34 class 3, and LY34/VU602 class 2 have been deposited in the PDB under accession codes 8TR2, 8TQB, 8TRD, 8TR0, and 8TRC, respectively, and in the Electron Microscopy Data Bank under accession codes EMD-41568, EMD-41501, EMD-41578, EMD-41567, and EMD-41577 respectively. They will be publicly available upon publication (HPUB status).

## Acknowledgments

We thank Drs. David Eliezer, Changhao He, and Nandish Khanra for helpful discussions. We thank all Levitz Lab members, especially Iram Arefin, for experimental assistance. We also thank Dr. Carl Fluck and Dr. Devrim Acehan at the Weill Cornell Medicine Cryo-EM Core facility for assistance with sample preparation as well as the Weill Cornell Scientific Computing Unit for their resources. Additionally, we are grateful to Drs. William Rice, Bing Wang, Huihui Kuang, and Alice Paquette at the NYU Langone Health’s Cryo–Electron Microscopy Laboratory for their assistance with data collection. We thank Dr. Asuka Inoue for the ΔGRK2/3 cells. We thank Dr. Craig Lindsley, Director of the Warren Center for Neuroscience Drug Discovery for his support. This work is supported by NIH grants F31NS129320 (A.S.), F32GM148001 (D.M.), and R01NS129904 (J. Levitz), the Margarita Salas Fellowship from the Ministry of Universities of Spain (A.G.-H), an NSF Graduate Research Fellowship (N.A), the Rohr Family Research Scholar Award (J. Levitz), and the Monique Weill-Caulier Award (J. Levitz).

## Author Contributions

Protein purification, Cryo-EM, and structural analysis were performed by A.S. with contributions from P.S., D.M., J.M., L.S., M.F. and J. Levitz. Live cell functional and conformational experiments were designed, performed, and analyzed by A.G.-H., J. Lee, N.A, M.K., and J. Levitz. K.G. and B.J.M. synthesized and characterized VU602. L.S. and M.F. designed and performed computational analysis. A.S. and J. Levitz wrote the paper with feedback from all authors.

## Declaration of Interests

The authors declare no conflicts of interest.

## Notes

https://wcm.box.com/s/810jehpv3ihgwj4rcfspa2a5t5cedgl9

### Competing Interest Statement

The authors have declared no competing interest.

