## Supplemental Information for "Structural basis of allosteric modulation of metabotropic glutamate receptor activation and desensitization"

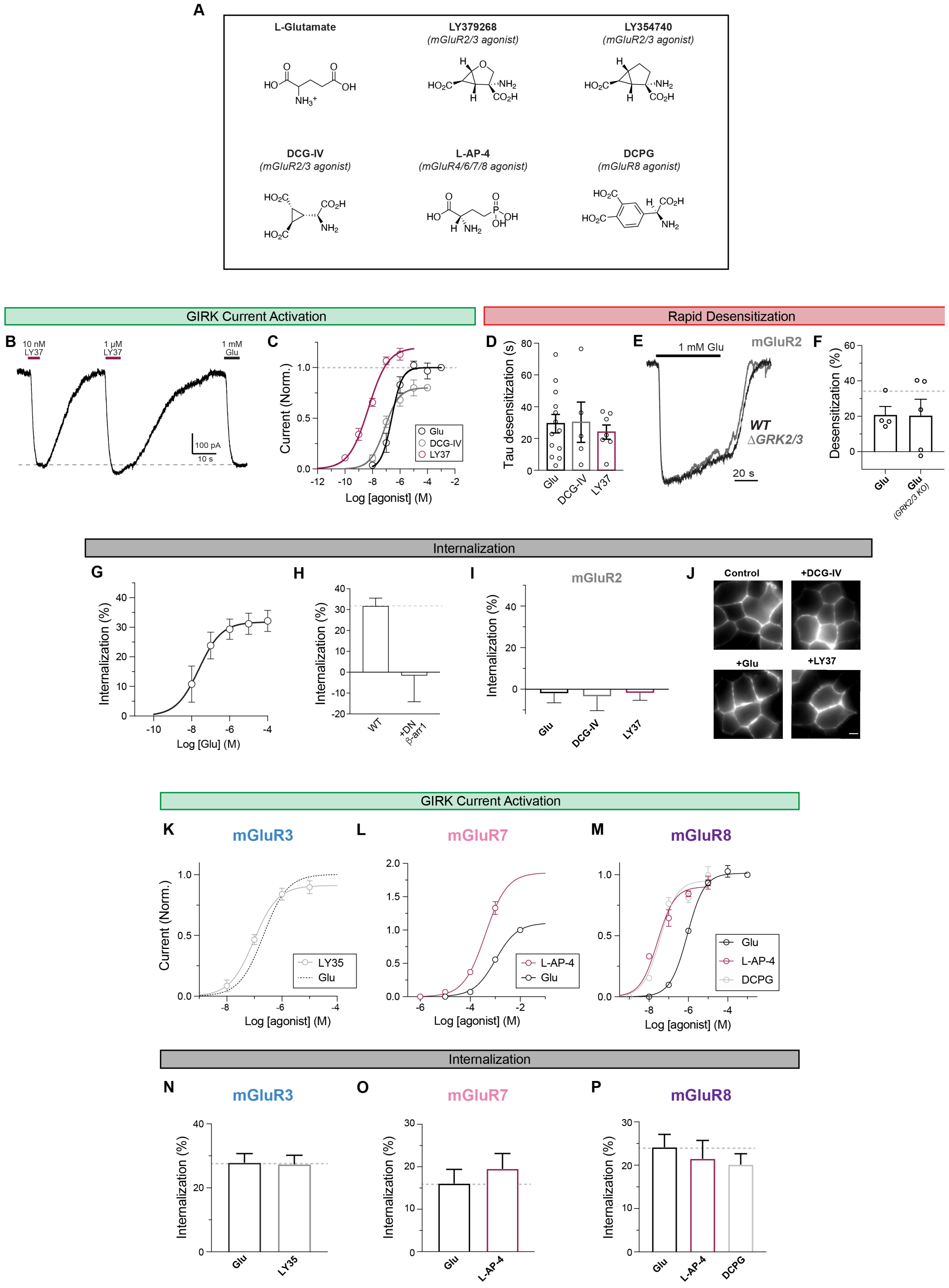
A
L-Glutamate
LY379268
(mGluR2/3 agonist)
LY354740
(mGluR2/3 agonist)
DCG-IV
(mGluR2/3 agonist)
L-AP-4
(mGluR4/6/7/8 agonist)
DCPG
(mGluR8 agonist)
GIRK Current Activation
Rapid Desensitization
B
10 nM LY37
1 μM LY37
1 mM Glu
100 pA
10 s
C
Current (Norm.)
Log [agonist] (M)
Glu
DCG-IV
LY37
D
Tau desensitization (s)
Glu
DCG-IV
LY37
E
1 mM Glu
mGluR2
WT
ΔGRK2/3
20 s
F
Desensitization (%)
Glu
Glu (GRK2/3 KO)
Internalization
G
Internalization (%)
Log [Glu] (M)
H
WT
+DN β-arr1
I
mGluR2
Glu
DCG-IV
LY37
J
Control
+DCG-IV
+Glu
+LY37
GIRK Current Activation
K
mGluR3
LY35
Glu
L
mGluR7
L-AP-4
Glu
M
mGluR8
Glu
L-AP-4
DCPG
Internalization
N
mGluR3
Glu
LY35
O
mGluR7
Glu
L-AP-4
P
mGluR8
Glu
L-AP-4
DCPG

**Figure S1. Further characterization of agonist driven G-protein activation and GRK/$\beta$-arr mediated desensitization across different mGluR subtypes.**

**(A)** Chemical structures for all agonists used in this study.

**(B)** Representative GIRK current trace showing dose-dependent responses to LY37 compared to glutamate.

**(C)** GIRK current dose-response curve for DCG-IV, glutamate and LY37 normalized to saturating glutamate. $EC_{50}$ = 84.9 ± 12.8 nM (DCG-IV); 206 ± 30 nM (glu); 6.3 ± 1.0 nM (LY37). n = 3-7 cells for concentration and agonist.

**(D)** Bar plot summarizing the kinetics of acute desensitization for the three agonists. Mean ± SEM, n= 5-15 cells.

**(E)** Representative traces of 1 mM glutamate induced acute desensitization for mGluR2 in WT HEK293 vs GRK2/3 KO HEK293 cells.

**(F)** Quantification of desensitization percentage at the end of 60 seconds of agonist application for glutamate in mGluR2. Grey dashed line represents mGluR3 mean desensitization to 100 µM Glu in equivalent conditions for WT cells (Fig. 1F).

**(G)** mGluR3 internalization dose-response curve for glutamate.

**(H)** Bar plot showing the internalization percentage of saturating glutamate for WT vs cells transfected with a dominant negative variant of $\beta$-arrestin (S412D).

**(I)** mGluR2 internalization percentage for all the three agonists in comparison to antagonist condition.

**(J)** Representative images of labelled SNAP-mGluR2 receptors under different agonists incubation for 30 min showing negligible internalization for this receptor.

**(K)** Dose-response curve of normalized current for LY35 agonist on mGluR3 in comparison with glutamate (dashed line). $EC_{50}$ = 95 ± 37 nM.

**(L)** Dose-response curve of normalized current for glutamate and L-AP4 agonists on mGluR7. $EC_{50}$ = 103 ± 26 µM (glu); 40 ± 18 µM (L-AP-4).

**(M)** Dose-response curves of normalized current for glutamate, L-AP4 and DCPG agonists on mGluR8. $EC_{50}$=0.9 ± 0.1 µM (glu); 27 ± 17 nM (L-AP-4); 36 ± 18 nM (DCPG)

**(N)** Internalization (%) caused by LY35 (10 uM) in comparison with glutamate (100µM) (mean represented by gray dashed line) on mGluR3

**(O)** Internalization (%) produced by L-AP4 (1 mM) and glutamate (10 mM) on mGluR7.

**(P)** Internalization (%) induced by glutamate (1 mM), L-AP4 (10 uM) and DCPG (10 uM) on mGluR8.

Bar-plots represent mean ± SEM for n>3 cells for electrophysiology experiments and n>30 images per condition (for at least 3 separate days) for imaging experiments.

For C, H and O, One-way ANOVA with multiple comparisons, n.s. p ≥ 0.05. For E, G and N, t-test; ** p<0.01; n.s. p≥0.05.

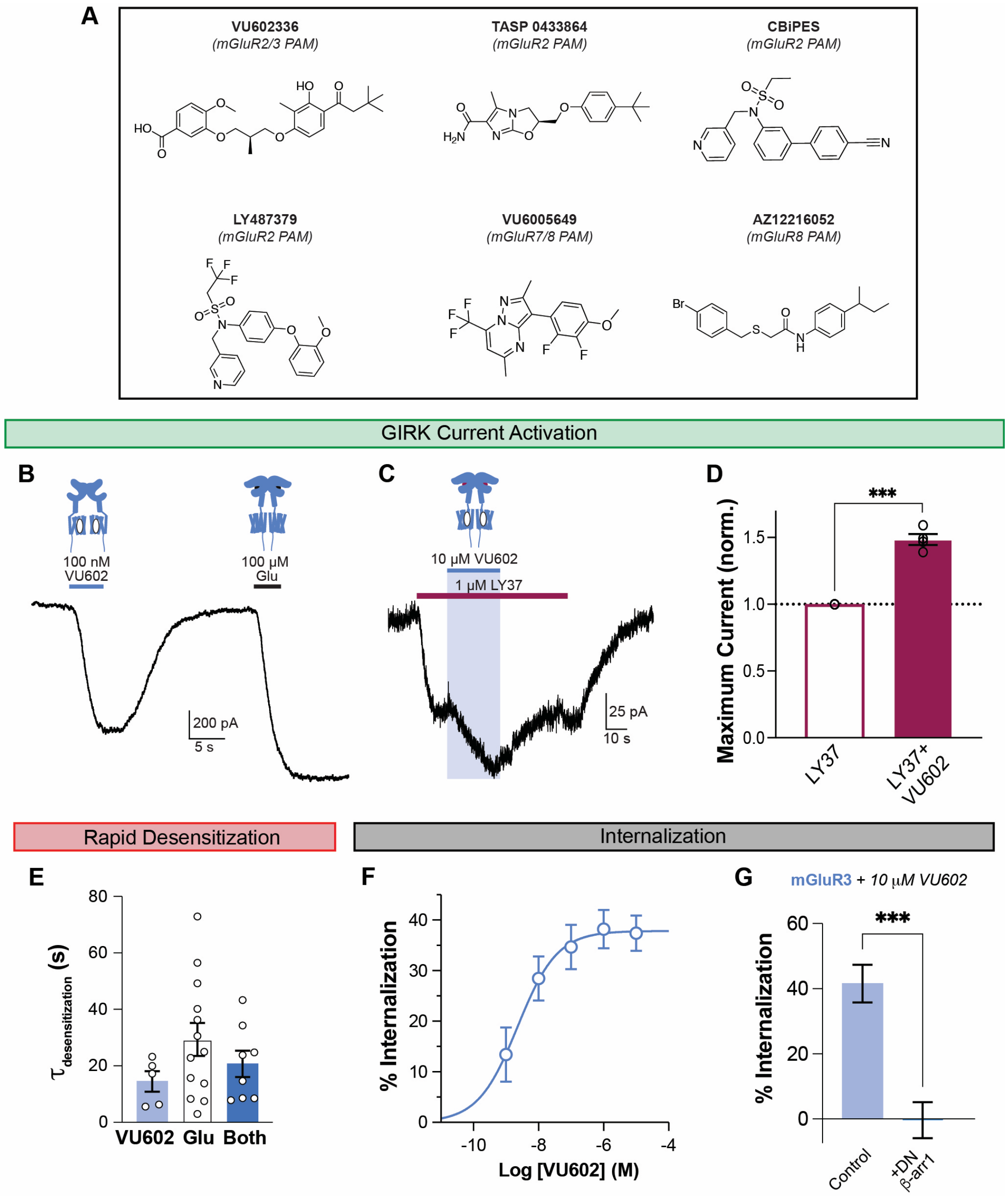

**Figure S2. Further analysis of the effects of VU602 PAM on mGluR3.**
**(A)** Chemical structures of the positive allosteric modulators used in this study.
**(B)** Representative trace of GIRK currents elicited by a sub-saturating dose of VU602 (100 nM) versus glutamate (100 μM).
**(C)** Representative trace showing that application of VU602 (10 μM) in the presence of saturating LY37 (1 μM) enhances inward GIRK currents.
**(D)** Quantification of the experiment in (C) normalized to the maximum amplitude of the agonist LY37 alone.
**(E)** Bar plot showing the tau of desensitization for the experiments in Fig. 2E and F.
**(F)** Internalization dose-response curve for VU602.

**(G)** Bar plot showing internalization (%) at saturating dose of VU602 comparing the control condition vs dominant negative b-arr1 cotransfection.
Bar-plots represent mean ± SEM for n>3 cells for electrophysiology experiments and n>30 images per condition (for at least 2 separate days) for imaging experiments.
For D and G, t-test, *** $p<0.001$. For E, One-way ANOVA with multiple comparisons; n.s. $p \geq 0.05$.

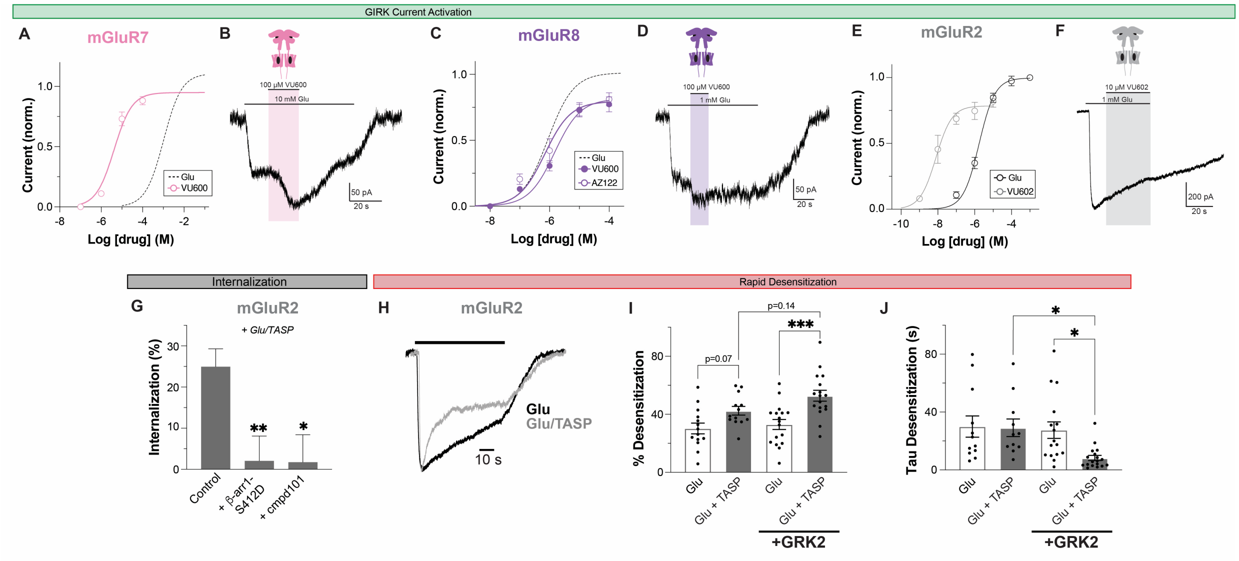

**Figure S3. Further characterization of PAM-driven G-protein activation and GRK/$\beta$-arr mediated desensitization across different mGluR subtypes.**

**(A)** VU600 dose-response curve on mGluR7 compared to glutamate (dashed line) using GIRK current assay. $EC_{50}$ = 4.2 ± 0.9 uM

**(B)** Representative trace showing that coapplication of VU600 on top of glutamate further potentiates GIRK activation by mGluR7.

**(C)** VU600 and AZ122 dose-response curves on mGluR8 compared to glutamate (dashed line) using GIRK current assay. $EC_{50}$= 1.5 ± 0.4 μM (VU600); 0.7 ± 0.3 μM (AZ)

**(D)** Representative trace showing that coapplication of VU600 on top of glutamate further potentiates GIRK activation by mGluR8.

**(E)** VU602 (gray) and glutamate (black) dose-response curves for mGluR2 using GIRK current assay. $EC_{50}$ = 7.9 ± 1.8 nM (VU602); 1.7 ± 0.2 μM (Glu)

**(F)** Representative trace showing that coapplication of VU602 on top of glutamate does not further potentiate GIRK activation by mGluR2.

**(G)** Internalization (%) observed in mGluR2 with the coapplication of PAM and glutamate is GRK and $\beta$-arr dependent as shown using the coexpression of a dominant negative $\beta$-arr1 or application of the GRK2/3 inhibitor cmpd101 (30 μM).

**(H)** Representative traces showing acute desensitization of mGluR2-mediated GIRK currents for glutamate or glutamate + TASP (PAM). Note: GRK2 is overexpressed in these cells to boost sensitivity.

**(I)** Quantification of the percentage of desensitization for glutamate or PAM + Glu in the control (first two bars; endogenous GRK expression) or with GRK2 overexpression (One-way ANOVA, *** $p<0.001$).

**(J)** Tau of desensitization in seconds for the experiments in H-I (One-way ANOVA, * $p<0.05$).

Bar-plots represent mean ± SEM for n>3 cells for electrophysiology experiments and n>30 images per condition (for at least 3 separate days) for imaging experiments.

For panels G, I and J, One-way ANOVA with multiple comparisons; * $p<0.05$,** $p<0.01$ and *** $p<0.001$.

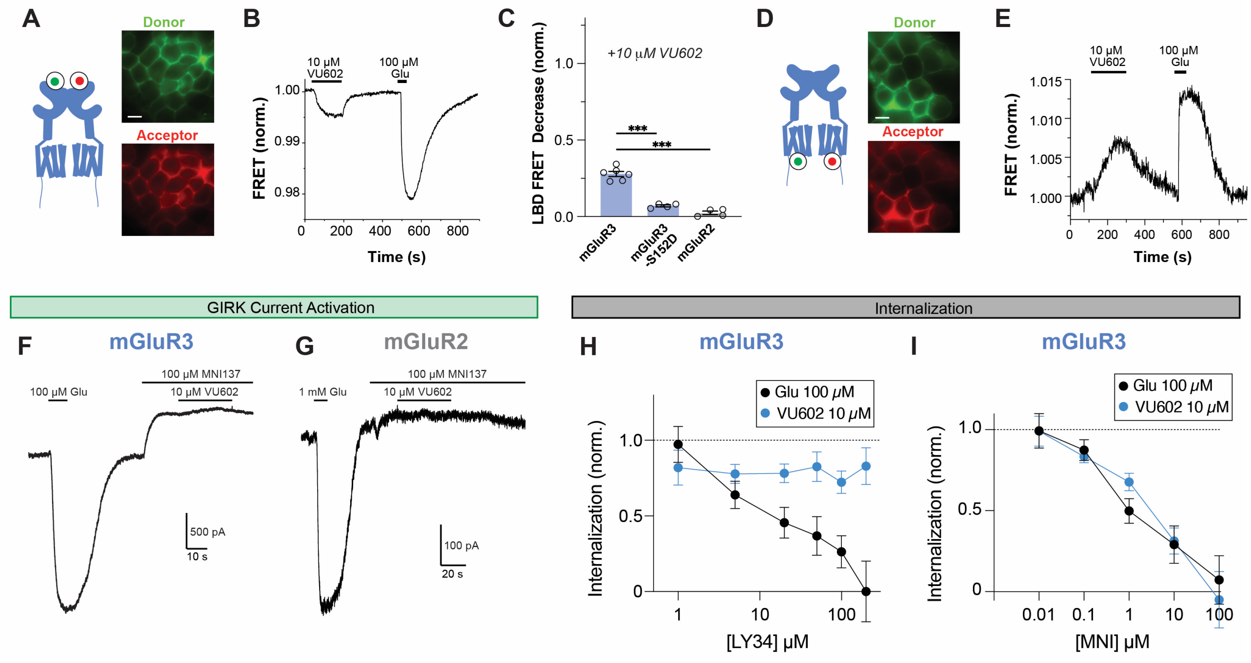

**Figure S4. Further analysis of the conformational rearrangements and functional effects of PAMs in combination with antagonists or NAMs.**

**(A)** Schematic (left) and representative images showing cells expressing SNAP-mGluR3 inter-LBD FRET sensor and labeled with donor and acceptor fluorophores. Scale bar = 10 $\mu$m.

**(B)** Representative inter-LBD FRET trace showing the response to saturating PAM and saturating glutamate.

**(C)** Summary bar graph showing the relative PAM response for wild type mGluR3, mGluR2, and mGluR3-S152D.

**(D-E)** Same as (A-B) but for inter-TMD FRET sensor.

**(F-G)** Representative traces showing that PAM-driven GIRK currents by mGluR3 (F) or mGluR2 (G) are fully blocked by preapplication of the NAM MNI-137.

**(H)** Titration of the antagonist LY34 has differential effects on mGluR3 internalization driven by saturating glutamate or saturating PAM. Values are plotted normalized to the amount of internalization seen in the absence of LY34.

**(I)** Titration of the NAM MNI-137 has strong effects on mGluR3 internalization driven by saturating glutamate or saturating PAM. Values are plotted normalized to the amount of internalization seen in the absence of MNI-137.

Dose-response curves in **H** and **I** represent mean ± SEM for each concentration for n>30 images per condition (for at least 3 separate days).

For panel C, One-way ANOVA with multiple comparisons. ***, $p<0.001$.

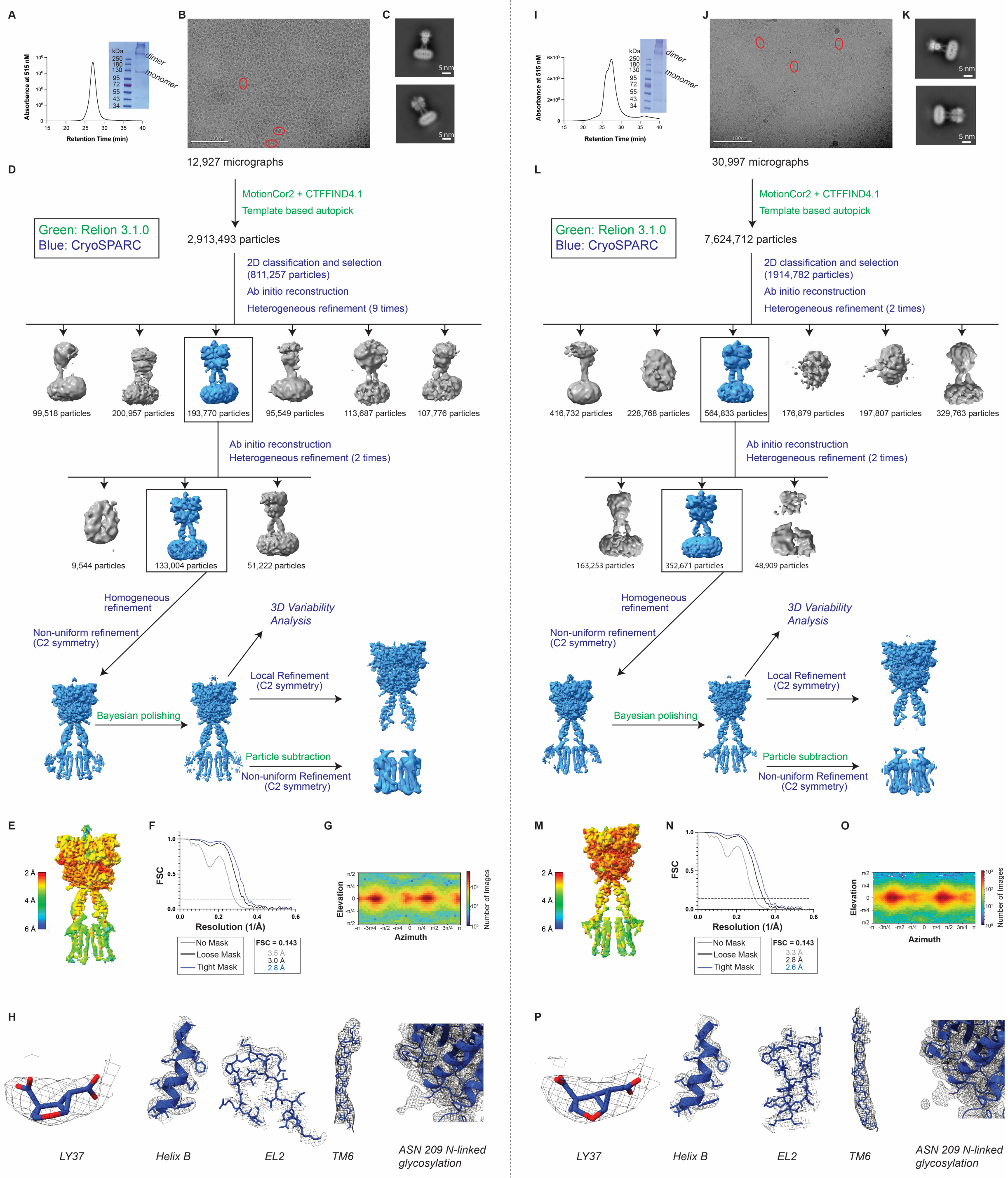

**Figure S5. Data processing workflow for agonist bound mGluR3 cryo-EM structures.**
**(A)** Size exclusion chromatography profile for mGluR3/LY37 data set with SDS PAGE gel of sample in inset.
**(B)** Representative mGluR3/LY37 micrograph with example particles highlighted.
**(C)** Representative 2D class averages of two different side views of mGluR3/LY37. Scale bar = 5 nm.

**(D)** Cryo-EM processing workflow for mGluR3/LY37. Maps are color coded as blue if particles are retained or gray if particles are discarded. Processing steps are colored green when performed in Relion version 3.1.0 and blue when performed in CryoSPARC version 3.3.2.
**(E)** Full-length cryo-EM density map colored by resolution as calculated in CryoSPARC.
**(F)** Fourier shell correlation (FSC) curves for LY37 bound mGluR3 density maps reported by CryoSPARC.
**(G)** Angular distribution heatmaps of particles in reconstructed LY37 bound mGluR3 map calculated in CryoSPARC.
**(H)** Select EM density and model. For LY37 and helix B (residues 102-115), the locally refined ECD map is shown at a threshold of 0.35 and zoned at 2 Å. For EL2, the full-length map density is shown at a threshold of 0.22 and zoned at 2 Å. For TM6, the TMD density resulting from particle subtraction is used at a threshold of 0.65 and zoned at 3 Å. To depict density assigned as N-linked glycosylation, the locally refined ECD map is shown at a threshold of 0.35 and zoned at 5.11 Å.
**(I-P)** Same as A-H for mGluR3/LY37+VU602 data set.

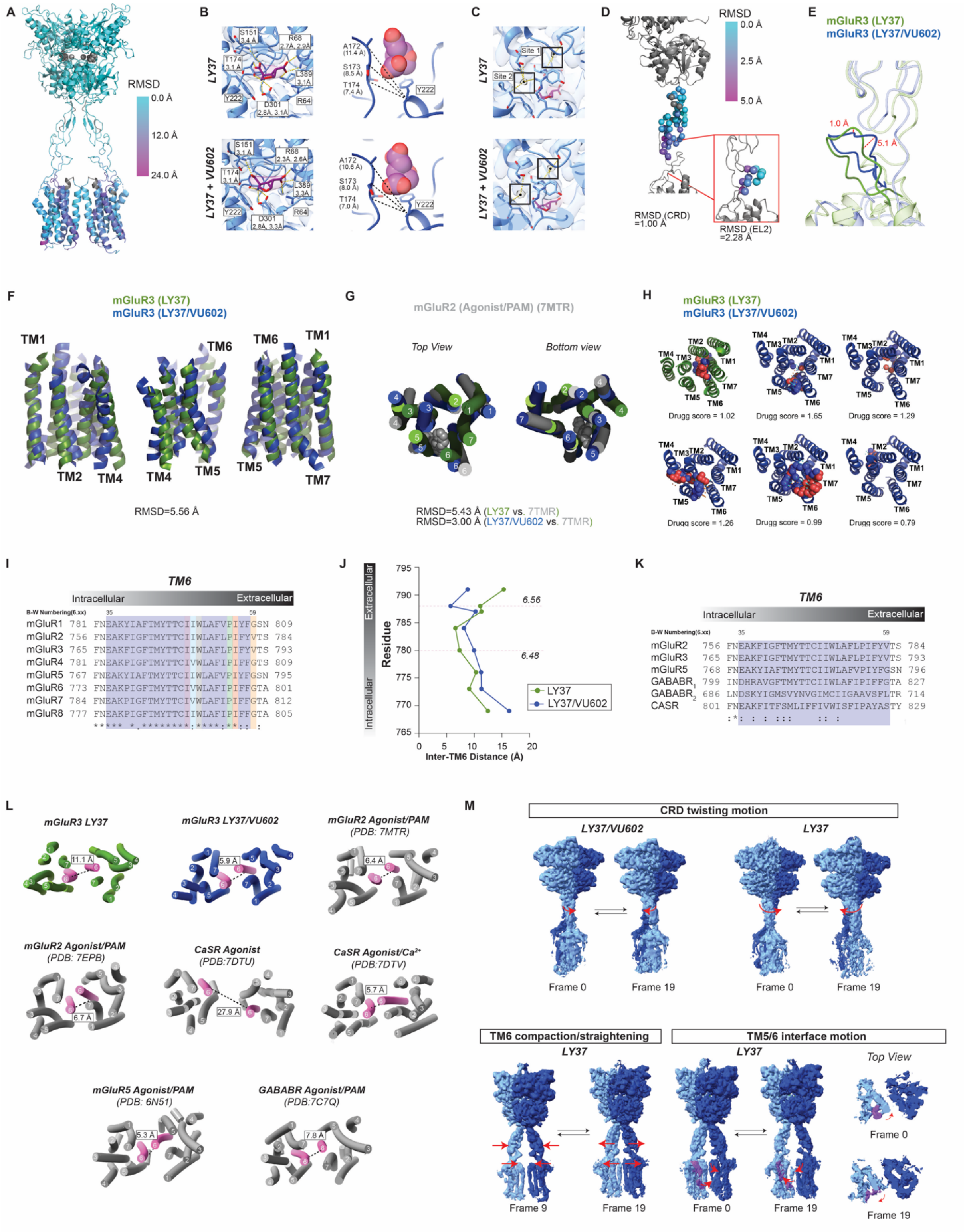

**Figure S6. Further comparative structural and functional analysis of agonist bound mGluRs.**
**(A)** Full length model of mGluR3 LY37/VU602 colored for RMSD calculated between the LY37 and LY37/VU602 models.

**(B)** Model, density, and stabilizing interactions for LY37 in the ligand binding domain (left). Measurements between top lobe residues and the bottom lobe residue Y222 (right).
**(C)** Two sites within the LBD where density assigned to ions is observed. mGluR3 is shown in blue, LY37 in purple, and ions in black.
**(D)** mGluR3 LY37/VU602 CRD colored by RMSD between mGluR3 LY37 and mGluR3 LY37/VU602 structures when aligned via the CRD of a single protomer or the EL2 of a single protomer (inset) with spheres showing C$\alpha$ position.
**(E)** EL2 offset when aligned via the CRD with distance change between C$\alpha s$ of K722 of the two models and E724 of the two models
**(F)** Side views comparing aligned TMDs for mGluR3 LY37 and mGluR3 LY37/VU602 structures.
**(G)** mGluR3 LY37, mGluR3 LY37/VU602 and mGluR2 (PDB: 7MTR) single TMD alignment with the mGluR2 ligand ADX55164 in gray spheres. Alignments are performed for the entire TMD, including loops.
**(H)** Top view of predicted druggable cavities in the TMDs of the mGluR3 LY37 and mGluR3 LY37/VU602 structures. Predicted cavities are depicted as spheres and colored according to their local properties (hydrophobic, cyan; aromatic, orange; hydrogen-bond acceptor and negative ionizable, blue; hydrogen-bond acceptor and positively ionizable, red).
**(I)** Sequence alignment of TM6 across all mGluR subtypes with Ballesteros–Weinstein numbering indicated.
**(J)** inter-TM6 interface profile measured by intersubunit C$\alpha$ distance of outward facing residues for LY37 and LY37/VU602 structures.
**(K)** TM6 sequence alignment for a variety of family C GPCRs.
**(L)** Top view of additional agonist bound class C GPCRs with distance measured from the C$\alpha$s of residue 6.56. TM6 is highlighted in pink.
**(M)** Single frames from 3D variability analysis with top row highlighting CRD twisting motion observed in both data sets and bottom row highlighting TM6 and CRD compaction as well as TM5/6 interface motion observed only in the mGluR3 LY37 data. See also supplemental videos 1-4.

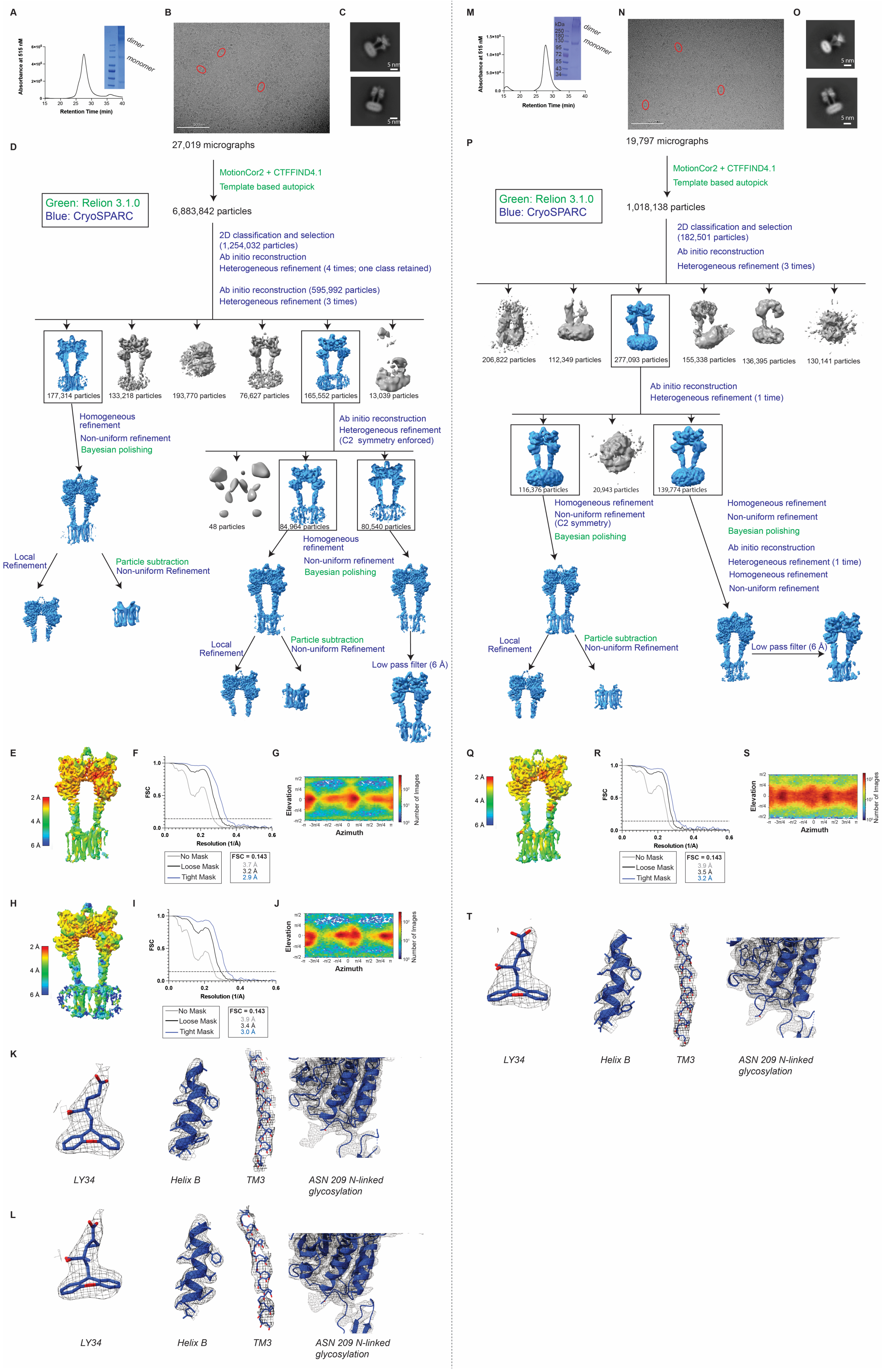

A
Absorbance at 515 nM
Retention Time (min)
dimer
monomer
B
C
5 nm
27,019 micrographs
D
MotionCor2 + CTFFIND4.1
Template based autopick
Green: Relion 3.1.0
Blue: CryoSPARC
6,883,842 particles
2D classification and selection
(1,254,032 particles)
Ab initio reconstruction
Heterogeneous refinement (4 times; one class retained)
Ab initio reconstruction (595,992 particles)
Heterogeneous refinement (3 times)
177,314 particles
133,218 particles
193,770 particles
76,627 particles
165,552 particles
13,039 particles
Homogeneous refinement
Non-uniform refinement
Bayesian polishing
Ab initio reconstruction
Heterogeneous refinement
(C2 symmetry enforced)
48 particles
84,964 particles
80,540 particles
Local Refinement
Particle subtraction
Non-uniform Refinement
Homogeneous refinement
Non-uniform refinement
Bayesian polishing
Low pass filter (6 Å)
E
F
G
2 Å
4 Å
6 Å
FSC
Resolution (1/Å)
No Mask
Loose Mask
Tight Mask
FSC = 0.143
3.7 Å
3.2 Å
2.9 Å
Elevation
Azimuth
Number of Images
H
I
J
3.9 Å
3.4 Å
3.0 Å
K
LY34
Helix B
TM3
ASN 209 N-linked glycosylation
L
LY34
Helix B
TM3
ASN 209 N-linked glycosylation
M
N
O
Absorbance at 515 nM
Retention Time (min)
kDa
250
180
130
95
72
55
43
34
dimer
monomer
19,797 micrographs
P
MotionCor2 + CTFFIND4.1
Template based autopick
Green: Relion 3.1.0
Blue: CryoSPARC
1,018,138 particles
2D classification and selection
(182,501 particles)
Ab initio reconstruction
Heterogeneous refinement (3 times)
206,822 particles
112,349 particles
277,093 particles
155,338 particles
136,395 particles
130,141 particles
Ab initio reconstruction
Heterogeneous refinement (1 time)
116,376 particles
20,943 particles
139,774 particles
Homogeneous refinement
Non-uniform refinement
(C2 symmetry)
Bayesian polishing
Homogeneous refinement
Non-uniform refinement
Bayesian polishing
Ab initio reconstruction
Heterogeneous refinement (1 time)
Homogeneous refinement
Non-uniform refinement
Local Refinement
Particle subtraction
Non-uniform Refinement
Low pass filter (6 Å)
Q
R
S
2 Å
4 Å
6 Å
FSC
Resolution (1/Å)
No Mask
Loose Mask
Tight Mask
FSC = 0.143
3.9 Å
3.5 Å
3.2 Å
Elevation
Azimuth
Number of Images
T
LY34
Helix B
TM3
ASN 209 N-linked glycosylation

**Figure S7. Data processing workflow for antagonist bound mGluR3 cryo-EM structures.**
**(A)** Size exclusion chromatography profile for mGluR3/LY34 data set with SDS PAGE gel of sample in inset.
**(B)** Representative mGluR3/LY34 micrograph with example particles highlighted.
**(C)** Representative 2D class averages of two different side views of mGluR3/LY34. Scale bar = 5 nm.
**(D)** Cryo-EM processing workflow for mGluR3/LY34. Maps are color coded as blue if particles are retained or gray if particles are discarded. Processing steps are colored green when performed in Relion version 3.1.0 and blue when performed in CryoSPARC version 3.3.2.
**(E)** Full length cryo-EM density map for LY34 class 3 colored by resolution as calculated in CryoSPARC.
**(F)** Fourier shell correlation (FSC) curves for LY34 class 3 bound density maps reported by CryoSPARC.
**(G)** Angular distribution heatmaps of particles in reconstructed LY34 class 3 map calculated in CryoSPARC.
(**H**) Full length cryo-EM density map for LY34 class 1 colored by resolution as calculated in CryoSPARC.
**(I)** Fourier shell correlation (FSC) curves for class 1 LY34 bound density maps reported by CryoSPARC.
**(J)** Angular distribution heatmaps of particles in reconstructed LY34 class 1 map calculated in CryoSPARC.
**(K)** Select LY34 class 3 EM density and model. For LY34 and helix B (residues 102-115), the locally refined ECD map is shown at a threshold of 0.35 and zoned at 2 Å. For EL2 and TM3 the full-length map density is shown at a threshold of 0.28 and zoned at 2 Å. To depict density assigned as N-linked glycosylation, the locally refined ECD map is shown at a threshold of 0.35 and zoned at 5.11 Å.
**(L)** Select LY34 class 1 EM density and model. For LY34 and helix B (residues 102-115), the locally refined ECD map is shown at a threshold of 0.35 and zoned at 2 Å. For EL2 and TM3 the full-length map density is shown at a threshold of 0.28 and zoned at 2 Å. To depict density assigned as N-linked glycosylation, the locally refined ECD map is shown at a threshold of 0.35 and zoned at 5.11 Å.
**(M-T)** Same as A-K for mGluR3/LY34+VU602 data set.

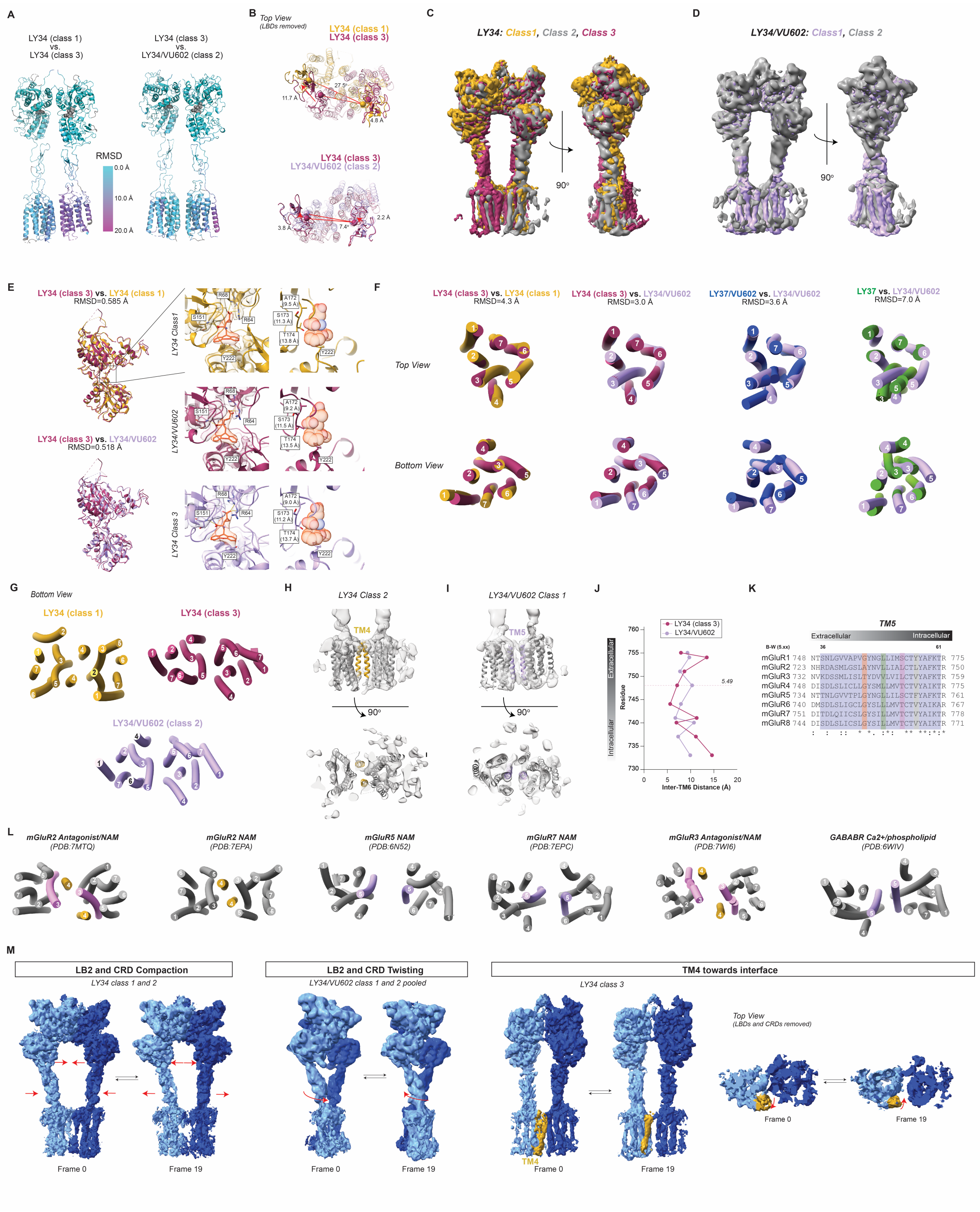

**Figure S8. Further comparative structural and functional analysis of antagonist bound mGluRs.**

**(A)** Full-length model of mGluR3 LY34 class 3 colored for RMSD calculated between the LY34 class 1 and class 3 models (left) or LY34 class 3 and LY34/VU602 class 2 models.

**(B)** Top view of LY34 class 1 (gold), LY34 class 3 (raspberry), and LY34/VU602 (lavender) when aligned via LB1 of chain A to highlight CRD offset.
**(C)** Aligned cryo-EM density maps for LY34 class 1 (gold), class 2 (grey), and class 3 (raspberry).
**(D)** Aligned cryo-EM density maps for LY34/VU602 class 1 (grey) and LY34/VU602 class 2 (lavender).
**(E)** Single LBD alignments for LY34 class 3 and LY34 class 1 or LY34 class3 and LY34/VU602 class 2 (top). Model, density, and stabilizing interactions for LY34 in the ligand binding domain (bottom).
**(F)** Top and bottom views of single TMD alignments between mGluR3 structures.
**(G)** Bottom view of TMDs for LY34 class 1 (gold), LY34 class 3 (raspberry), and LY34/VU602 class 2 (lavender).
**(H)** Individual LY34 class 1 TMD bundles aligned to the LY34 class 2 cryo-EM density to identify a putative TM4 containing interface highlighted in gold.
**(I)** Individual LY34 class 3 TMD bundles aligned to the LY34/VU602 class 1 cryo-EM density to identify a putative TM5 containing interface hlighted in lavender.
**(J)** Inter-TM5 interface profile measured by intersubunit C$\alpha$ distance of outward facing residues for LY34 class 3 and LY34/VU602 class 2.
**(K)** Sequence alignment of TM5 across all mGluR subtypes with Ballesteros–Weinstein numbering indicated.
**(L)** Top view of additional antagonist and/or NAM -bound class C GPCRs with TM3 highlighted in pink, TM4 highlighted in gold, and TM 5 highlighted in lavender.
**(M)** Single frames from 3D variability analysis as depicted in supplemental videos 5-7. They highlight compaction and twisting motions at the CRD and LB2 interface as well as changes in TMD dimer interface in the LY34 class 3 data.

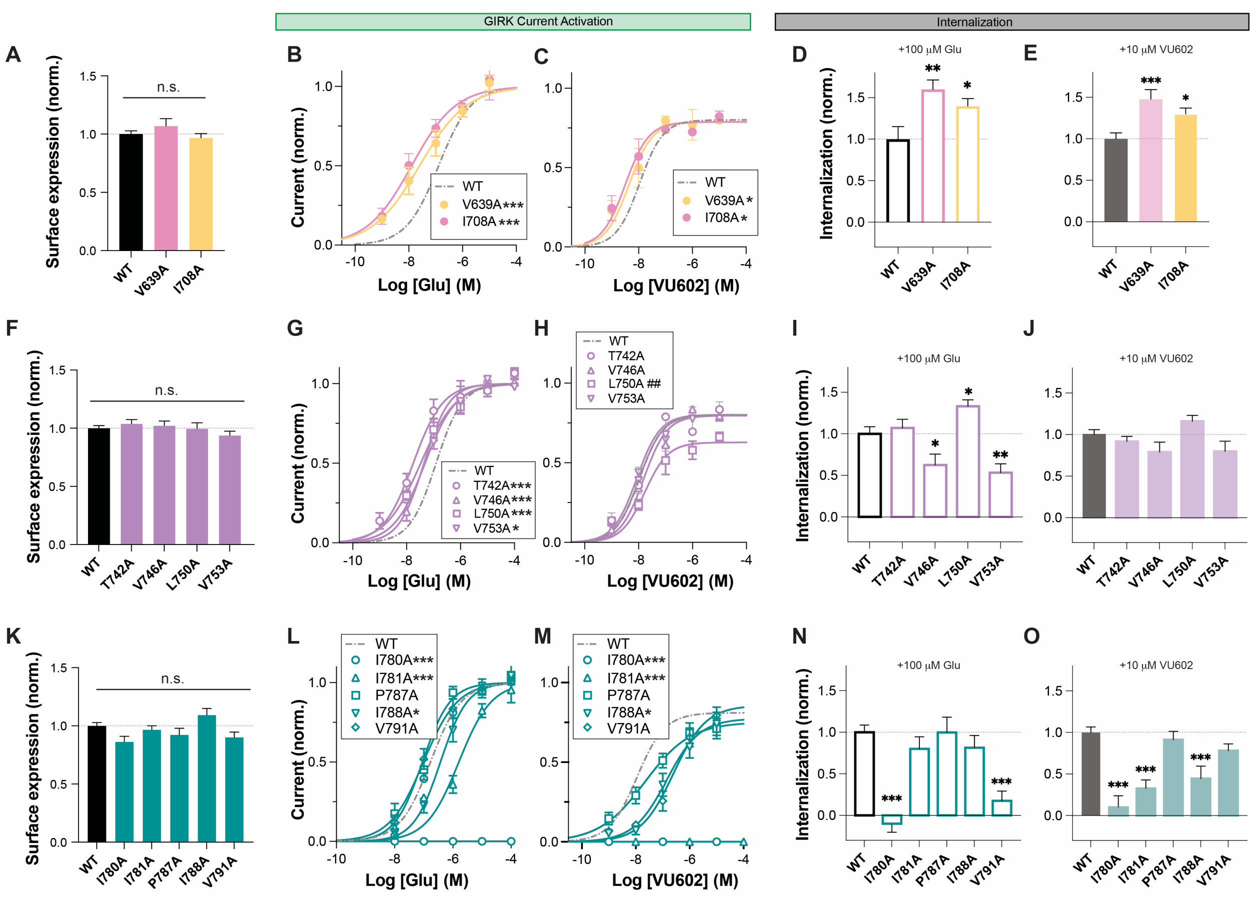

**Figure S9. Summary of alanine scanning mutagenesis analysis of inter-TMD interfaces**
**(A-E)** Evaluation of the effects of TM3/TM4 mGluR3 alanine mutants in G protein activation and internalization assays. (A) Surface expression quantification for the two mutants normalized to glutamate. (B-C) Normalized glutamate (B) and VU602 (C) GIRK current dose-response curves (dotted grey line represents WT). $EC_{50}$ values for these mutants are: (B) V639A: 20.8 ± 7 nM; I708A: 11.5 ± 3 nM and (C) V639: 2.8 ± 2.1 nM and I708A: 2.2 ± 1 nM for VU602. (D-E) Internalization produced by glutamate (D) or VU602 (E) normalized to WT.
**(F-O)** Same as A-E but for TM5 **(F-J)** and TM6 mutants **(K-O)**. $EC_{50}$ values are: **(G)** T742A: 16.9 ± 3 nM; V746A: 45.8 ± 8.8 nM; L750A: 30.9 ± 6.4 nM; V753A: 58.9 ± 6.8 nM; **(H)** T742A: 8.5 ± 1.3 nM; V746A: 15.3 ± 7.6 nM; L750A: 16.9 ± 7 nM; V753A: 10.5 ± 4.6 nM; **(L)** I781A: 1.9 ± 0.4 μM; P787A: 95.1 ± 16.8 nM; I788A: 378 ± 52.5 nM; V791A: 97.6 ± 16.5 nM; **(M)** P787A: 14.9 ± 8.6 nM; I788A: 109.9 ± 62 nM; V791A: 205.9 ± 91 nM.
Bar plots represent mean ± SEM for n>30 images per condition from at least 3 separate days. Dose response curves represent mean ± SEM for each glutamate or PAM concentration for at least 3 different cells per concentration.
For A, D, E, F, I, J, K, N and O, One-way ANOVA with multiple comparisons; *p<0.05, ***p<0.001. For B, C, G, H, L and M, $EC_{50}$ shift was evaluated using an F-test of each fitted curve $EC_{50}$ where the null hypothesis was “$EC_{50}$ ratio is shared among WT and mutants”. *p<0.05; ***p<0.001.

| Data collection and processing | LY37 | LY37/VU602 | LY34 (class 1) | LY34 (class3) | LY34/VU602 |
|---|---|---|---|---|---|
| Microscope | Krios | Krios | Krios | Krios | Krios |
| Magnification | 105,000X | 105,000X | 105,000X | 105,000X | 105,000X |
| Voltage (kV) | 300 | 300 | 300 | 300 | 300 |
| Electron exposure (e-/Å^2) | 59.18 and 56.84 | 57.42 and 57.84 and 58.11 and 58.45 | 57.07 and 58.80 | 57.07 and 58.81 | 52.26 |
| Defocus range (μm) | 1.2 – 1.7 and 1.3 – 2.0 | 0.9 - 1.9 and 0.7 - 2.0 | 0.8 – 2.4 and 0.7 - 2.0 | 0.8 – 2.4 and 0.7 - 2.1 | 0.8 - 2.4 and 0.5 - 2.8 |
| Pixel Size (Å) | 0.426 | 0.4125 | 0.4125 | 0.4125 | 0.4125 |
| Binned Pixel size (Å) | 0.852 | 0.825 | 0.825 | 0.825 | 0.825 |
| Frames/movies (#) | 40 | 50 | 50 | 50 | 60 and 45 |
| Symmetry imposed | C2 | C2 | C1 | C1 | C2 |
| Movies (#) | 19,797 | 30,997 | 27,019 | 27,019 | 19,797 |
| Particle images (#) | 133,004 | 352,671 | 84,964 | 117,314 | 116,376 |
| Box size (pixels) | 416 | 416 | 416 | 416 | 416 |
| Global map resolution (Å) | 3.5 | 3.3 | 3.4 | 3.2 | 3.3 |
| FSC threshold | 0.143 | 0.143 | 0.143 | 0.143 | 0.143 |
| **Refinement** | | | | | |
| Composition (#) | | | | | |
| Non-hydrogen atoms | 10112 | 10244 | 9622 | 10052 | 9856 |
| Protein residues | 1490 | 1488 | 1468 | 1488 | 1474 |
| Ligands | 6 | 6 | 2 | 2 | 2 |
| R.m.s deviations | | | | | |
| Length (Å) | 0.016 | 0.016 | 0.016 | 0.005 | 0.018 |
| Angles (°) | 1.092 | 1.138 | 1.387 | 0.929 | 1.387 |
| Validation | | | 2.35 | | |
| MolProbity score | 1.86 | 1.88 | 2.35 | 1.94 | 2.45 |
| Clash core | 6.10 | 5.98 | 8.74 | 7.66 | 13.77 |
| Rotamer outliers (%) | 0.22 | 1.28 | 1.94 | 0.56 | 1.50 |
| Ramachandran plot (%) | | | | | |
| Outliers | 0.21 | 0.00 | 0.00 | 0.00 | 0.21 |
| Allowed | 9.04 | 7.65 | 15.08 | 8.92 | 15.33 |
| Favored | 90.75 | 92.35 | 84.92 | 91.08 | 84.47 |

**Table S1: Cryo-EM data collection and refinement statistics**
Information on cryo-EM data collection settings and refinement statistics for all mGluR3 models.

### Supplemental Video Legends:

**Supplemental Video 1: Conformational variability in LY37 bound mGluR3 TM6 and CRD compaction**
3D variability analysis for the LY37 bound data revealing a compaction motion at the CRD and a compaction and straightening of the TM6 dimer interface.

**Supplemental Video 2: Conformational variability in LY37 bound mGluR3 TMD dimer interface**
3D variability analysis for the LY37 bound data showing a rotation at the TMD to bring TM5 towards the dimer interface.

**Supplemental Video 3: Conformational variability in LY37 bound mGluR3 CRD position**
3D variability analysis for the LY37 bound data showing a twisting of the CRD in which they pass each other with a large range of motion.

**Supplemental Video 4: Conformational variability in LY37/VU602 bound mGluR3 CRD position**
3D variability analysis for the LY37/VU602 bound data showing a twisting of the CRD in which they pass each other with a smaller range of motion compared to the LY37 only data (Supplemental Video 3).

**Supplemental Video 5: Conformational variability of the mGluR3 LBDs and CRDs in LY34 class 1 and 2**
3D variability analysis of changes in LB2 distance and CRD compaction for the pooled LY34 class 1 and class 2 data.

**Supplemental Video 6: Conformational variability of the mGluR3 LBDs and CRDs in LY34/VU602 class 1 and 2**
3D variability analysis of changes in LB2 distance and CRD compaction for the pooled LY34/VU602 class 1 and class 2 data.

**Supplemental Video 7: Conformational variability of TMD dimer interface in LY34 class 3**
3D variability analysis of changes in TMD so that TM4 faces towards the dimer interface for LY34 class 3 data only.

All supplemental videos are available for download:
https://wcm.box.com/s/810jehpv3ihgwj4rcfspa2a5t5cedgl9
